# Intravital imaging the formation and resolution of MHC class II-positive T cell activation niches

**DOI:** 10.1101/2025.08.05.668796

**Authors:** David Oleksyn, Jim Miller

**Affiliations:** Center for Vaccine Biology and Immunology and Department of Microbiology and Immunology University of Rochester Medical Center, Rochester, NY

**Keywords:** Chemokines, inflammation, Th1 cells, spectral unmixing, CD8 fluorescent reporter mice, two-photon microscopy

## Abstract

Intravital imaging has revealed many of the cellular interactions that regulate immune responses, but is limited by the number of cells that can be simultaneously identified and often restricted to analysis of a single time point. We have developed two new fluorescent reporter strains, IEbeta-mAmetrine, and CD8beta-LSSmOrange, that faithfully label cells expressing MHC class II and CD8-positive conventional T cells, respectively. These fluorescent proteins are spectrally distinct from commonly used fluorescent proteins (GFP, YFP) and so these mice can be used in combination with many previously created reporter mice. In addition, we established a protocol where we can sequentially image the same area of the ear dermis over several weeks without inducing any inflammation. We applied these techniques to IEbeta-mAmetrine mice co-expressing markers for CD11c, CXCL10, and CD4 T cells to quantify the formation of CXCL3 activation clusters, elaboration of different MHC class II positive cells within these clusters, accumulation of CD4 T cells within these clusters, and the dissipation of these T cell activation niches as the inflammatory response wanes.

**Summary Blurb:** This study describes the development and utility of two new fluorescent reporter mice for intravital imaging of the same tissue site to follow the kinetics of an immune response.

## Introduction

An effective immune response requires sequential interactions between different populations of migratory immune cells within lymphoid tissue and again at peripheral sites of inflammation and infection. Recent imaging experiments, in particular intravital imaging (IVM), have shown that many of these interactions take place within dynamically and spatially controlled cell clusters both within and without lymphoid tissues (Bala N et al, 2022). In the lymph node, chemokine driven migration and localization within discrete environments drive T cells activation and differentiation (Groom JR et al, 2012, Leal JM et al, 2021, Ugur M & Mueller SN, 2019). This is most elegantly illustrated in the dynamic choreography of cell migration during the formation of a germinal center (Victora GD & Nussenzweig MC, 2022). Although less well understood, similar localization and clustering of T cells can regulate T cell activation and impact T cell fate in peripheral tissues (Natsuaki Y et al, 2014, Prizant H et al, 2021) and tumors (Di Pilato M et al, 2021, Jansen CS et al, 2019).

The development of IVM has been instrumental in revealing the spatial and temporal dynamics of cell migrations that regulate and mediate immune responses REF.

One limitation to IVM is the number of specific cells types that can be simultaneously imaged. This is due in part to the availability of mice expressing cell type specific fluorescent proteins that can be spectrally distinguished. A second limitation to IVM is the ability to image the same tissue site in a single animal over the course of an entire immune response. As a result, many studies focus on a single time point often at the peak of the response.

In this report we have addressed these limitations by creating two new tissue specific reporter mouse models, expressing mAmetrine in MHC class II positive cells (IEbeta-mAmetrine) and expressing LSSmOrange in conventional CD8 positive cells (CD8beta-LSSmOrange). In addition, we have modified the existing dermal imaging protocol, so that we can image the same region of mouse skin dermis over the entire length of an inflammatory response. We applied these techniques to mice co-expressing markers for MHC class II, CD11c, CXCL10, and CD4 T cells to quantify the formation of CXCL3 activation clusters, elaboration of different MHC class II positive cells within these clusters, accumulation of CD4 T cells within these clusters, and the dissipation of these T cell activation niches (Prizant H et al, 2021) as the inflammatory response wanes.

## Results and Discussion

### IEbeta-mAmetrine and CD8beta-LSSmOrgage mice

To generate new animals for intraviral imaging for immune cell types, we first screened candidate fluorescent proteins for two photon excitation and spectral overlap. The goal was to identify fluorescent proteins that were spectrally distinct from commonly used fluorescent proteins (such as GFP and YFP, so that any animal created could be crossed to common existing strains. We settled on using mAmetrine and LSSmOrange (see supplemental Table S1)

MHC class II reporter strains have been previously generated by fusing GFP to the carboxyterminus of IAbeta and have been useful for tracking the intracellular localization and turnover of class II proteins (Bannard O et al, 2016, Chieppa M et al, 2006, Litingtung Y et al, 2002). But such fusion proteins have the potential to induce misregulation of the target protein. To generate a fluorescent marker strain that does not interfere with MHC class II protein structure, we took advantage that C57BL/6 mice maintain a fully functional MHC class II IEbeta gene, but do not express and IE protein due to a defective IEalpha gene. When these mice were reconstituted with a functional IEalpha gene, normal and functional cell surface expression of MHC class II IE was detected (Le Meur M et al, 1985, Pinkert CA et al, 1985, Yamamura K-i et al, 1985). Therefore, we chose to use CRISPR to knock the mAmetrine coding sequence (including a beta globin 3’UTR and polyA addition site) into the start site of IEbeta translation, conferring the genetic locus control of IE to mAmetrine (Fig 1 and supplemental Table S2). Although this construct knocks out IEbeta expression, it should have no impact on the expression of class II, as C57BL/6 mice do not normally express IE and should continue to express normal levels of endogenous MHC class II IA proteins.

**Figure 1.**
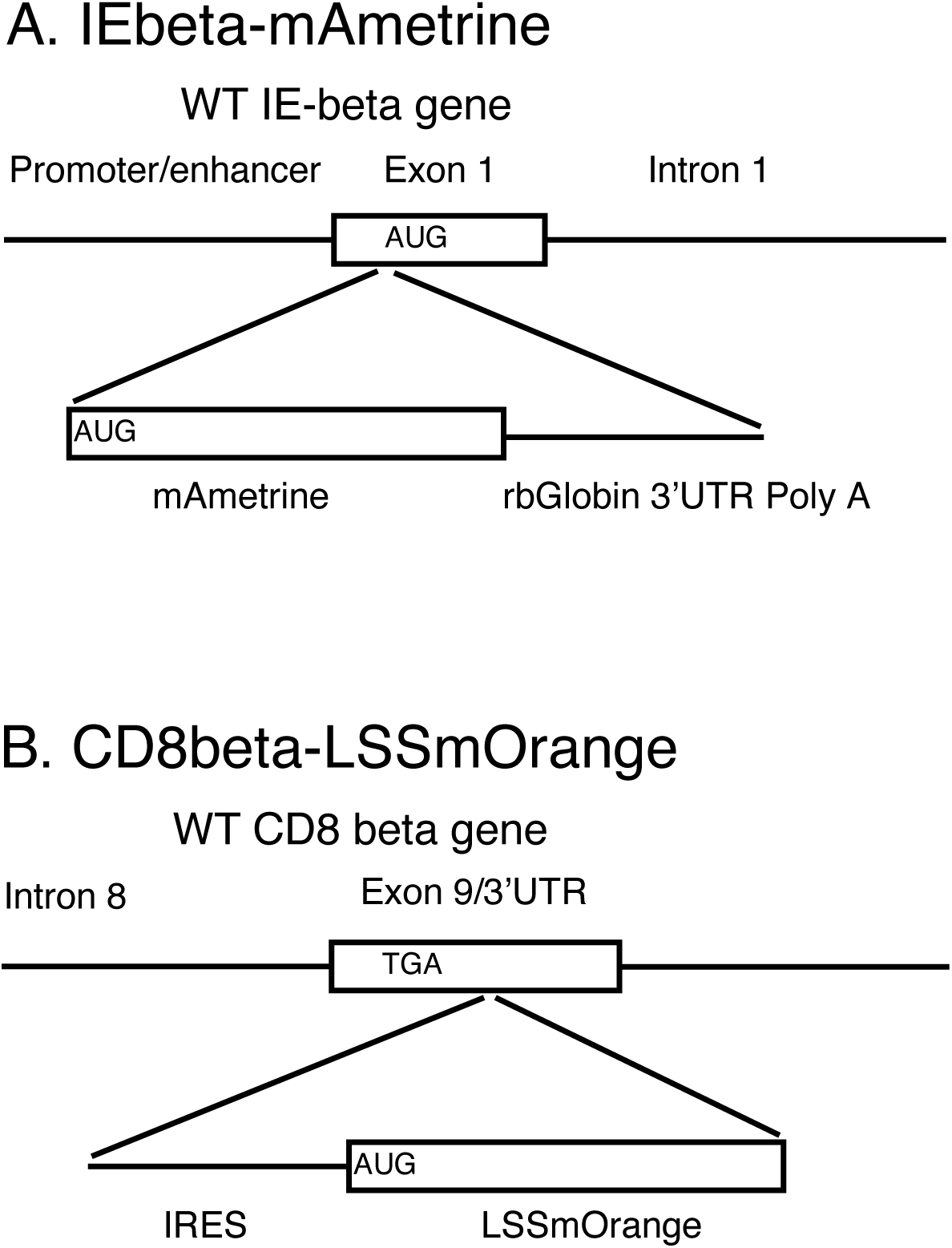
Generation of tissue specific fluorescent reporter mice. **(A)** The cassette containing the mAmetrine coding sequence and a rabbit beta globin 3’UTR/poly A addition site was inserted at the start site of the IE-beta gene. **(B)** A cassette containing an IRES and the LSSmOrange coding region was inserted into the 3’UTR shortly following the termination codon of the CD8-beta gene.

A reporter strain expressing TdTomato under control of the CD8alpha gene has been previously generated (Mohan JF et al, 2017). However, CD8alpha is not a T cell specific lineage marker and CD8alpha-alpha homodimers can be expressed on a variety of cell types, including dendritic cells, natural killer cells, interstitial epithelial lymphocytes, and gamma delta T cells (Gangadharan D & Cheroutre H, 2004, Srinivasan S et al, 2024). In contrast, CD8beta expression is highly restricted to conventional CD8 T cells. Therefore, we used CRISPR to knock in a cassette containing an IRES and the LSSmOrange coding region into the 3’UTR of the endogenous CD8beta gene (Fig 1 and supplemental Table S3). This construct will produce a bicistronic message that should allow for normal expression of CD8beta from the 5’ Kozak start sequence and LSSmOrange from the IRES in the same mRNA.

### Expression of IEbeta-mAmetrine

To determine whether expression of mAmetrine was a faithful reporter of MHC class II expression, we harvested cells from lymph nodes, spleen, and lungs from WT C57BL/6 mice and from homozygous IEbeta-mAmetrine mice and stained them for endogenous MHC class II IA^b^. As can be seen in Fig 2, there is good concordance between mAmetrine expression driven from the IEbeta knockin and endogenous IA^b^. In the total cell populations (Fig 2A) the majority of class II positive cells are B cells (Fig 2B). To highlight the myeloid class II positive cells, we gated on CD3- and CD19- cells. There is more heterogeneity in the levels of class II expression in this population, but again there is a good concordance between mAmetrine and IA^b^ (Fig 2C). Expression of mAmetrine did not impact the development of CD4 single positive cells in the thymus (supplemental Fig S1) or the relative number of different class II-positive antigen presenting populations (Fig 3A,B). Taking advantage of the different level of endogenous class II on these subpopulations in different tissue, we found a significant correlation between the level of endogenous IA^b^ and mAmetrine in the IEbeta-mAmetrine mice (Fig 3C). The correlation is not perfect likely due to differential half-lives of IA^b^ and mAmetrine proteins.

**Figure 2.**
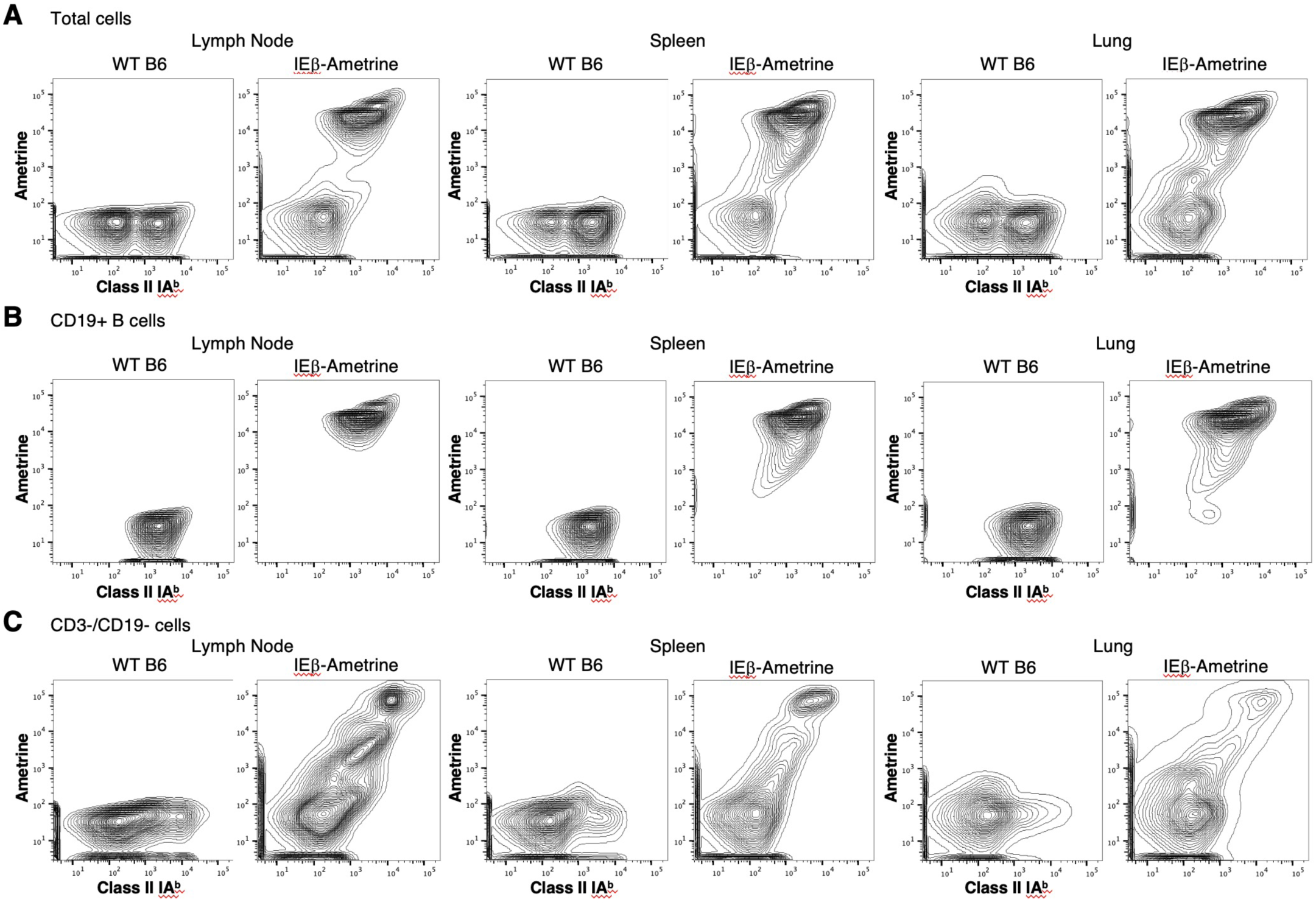
Expression of mAmetrine in the IEbeta-mAmetrine mice is a faithful reporter for MHC class II. Representative flow cytometry of cells isolated from lymph nodes, spleens, or lungs of WT or IEbeta- mAmertine mice and stained for endogenous class II I-A^b^ and displayed with Ametrine expression. **(A)** total cells from each tissue. **(B)** samples gated on CD19+ B cells. **(C)** samples gated on CD3-, CD19- cells.

**Figure 3.**
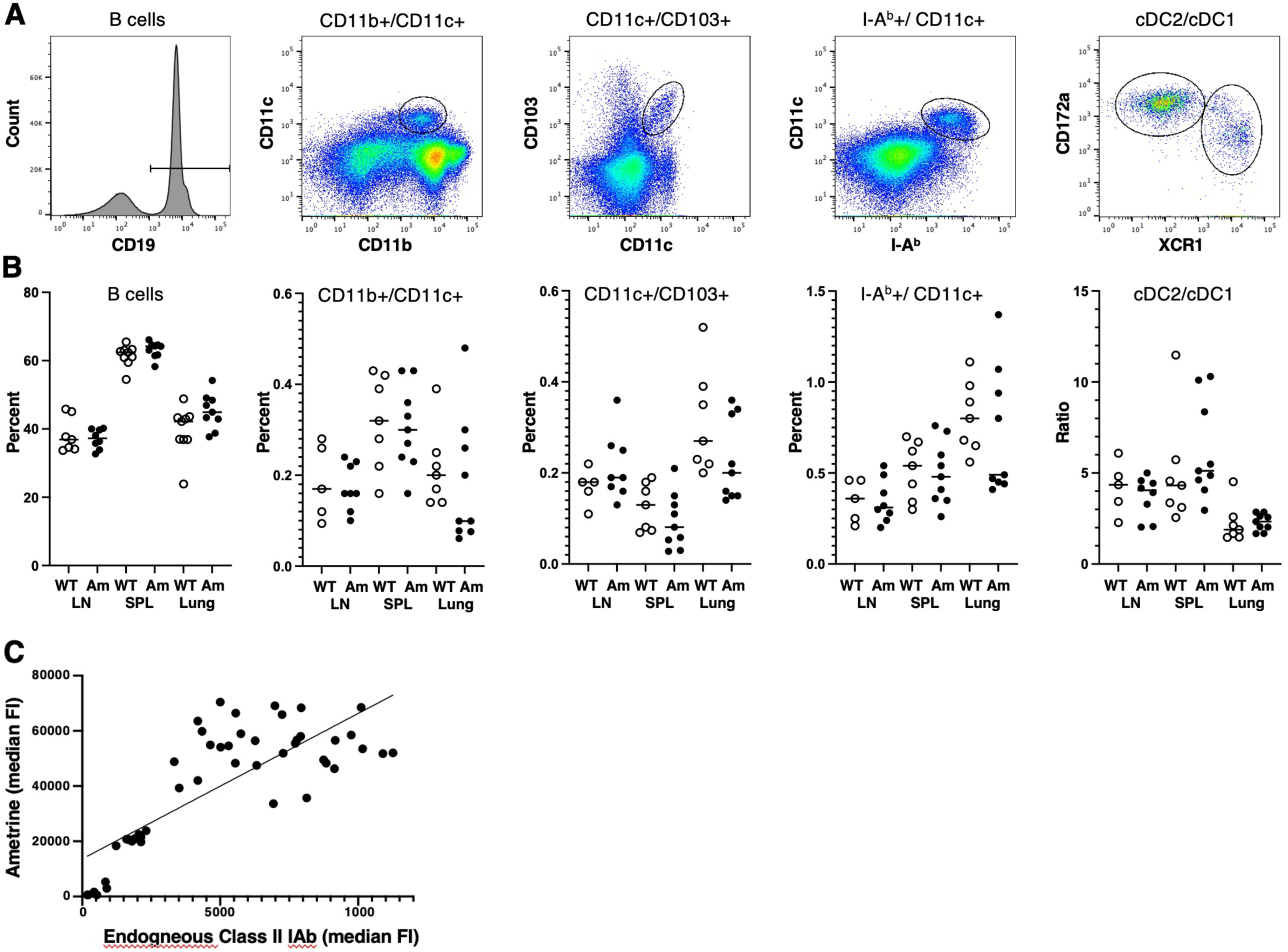
Expression of IEbeta-mAmetrine does not alter the frequency of antigen presenting cell populations. **(A)** Example of gating strategy to identify APC cell populations. B cells are shown for total population, other cells are pregated on CD19-/CD3- cells, and cDC1/cDC2 cells are further pregated on IA^b^+/CD11c+ cells. **(B)** Cells were isolated from lymph node (LN), spleen (SPL), and Lung of WT (open symbols) and IEbeta-mAmetrine (Am; filled symbols)) mice and the percentage of B cells, CD11b+/CD11c+, CD11c+/CD103+, and IA^b^+/CD11c+ cells (based on total cells isolated from the tissue) and the ratio of cDC2/cDC1 (subgated from IA^b^+/CD11c+ cells) are shown for 3 separate experiments containing 1-3 mice per group (n=5-9; no significant differences between WT and IEbeta-Ametrine for any of the populations). **(C)** Median fluorescent intensity of Ametrine by endogenous MHC class II IA^b^ expression in gated subpopulations (B cells, cDC1, cDC2, CD103 DC, CD11c+, CD11b+/CD11c+) from lymph node, spleen, and lung of IEbeta-Ametrine mice. (R^2^=0.62, p<0.0001) .

To assess the expression of IEbeta-mAmetrine within intact tissue, we imaged the epidermal layer of mouse ear skin by IVM (Fig 4A). The mice were maintained on an C57BL/6-Albino background to eliminate spectral interference with melanin (Sabino CP et al, 2016). Expression of mAmetrine clearly identified the MHC class II-positive Langerhans cells (LC) that reside above the collagen rich dermis and display distinct dendritic cell morphology as be seen in the higher magnification image (Fig 4A) (Romani N et al, 2010, Tong PL et al, 2015). To determine if mAmetrine can also be used to identify MHC class II-positive thymic epithelial cells, thymus was removed from IEbeta-mAmetrine mice and imaged ex vivo on the two-photon microscope. Cells expressing mAmetrine with epithelial cell morphology were clearly visible in both the cortical region of the thymus (Top image in Fig 4B) and deeper in the medullary region (bottom image in Fig 4B) (Yang SJ et al, 2006). Collectively the flow data and the tissue imaging data indicate that mAmetrine expression is a faithful reporter of MHC class II expression in IEbeta- mAmetrine mice.

**Figure 4.**
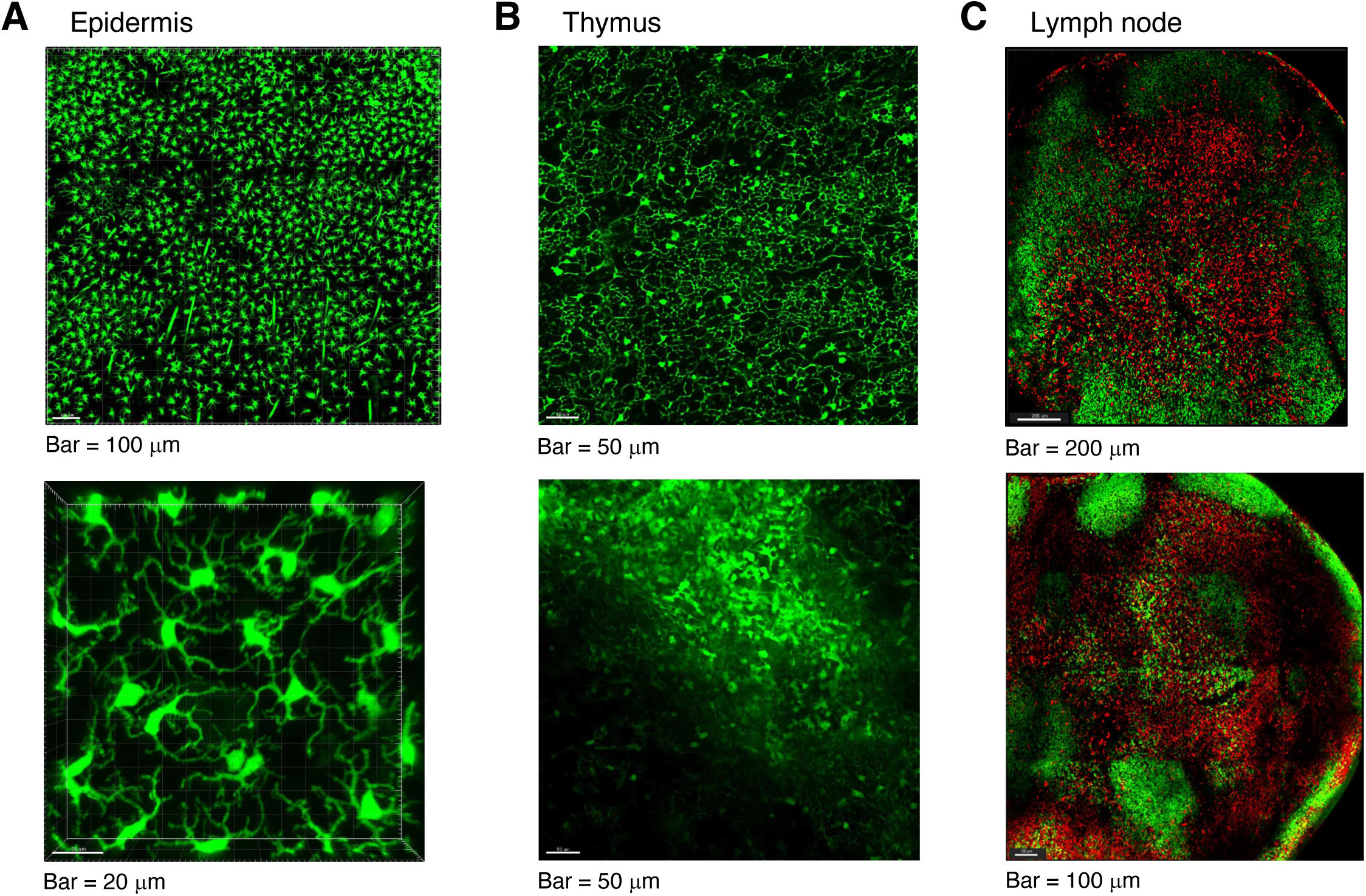
Expression of IEbeta-mAmetrine and CD8beta-LSSmOrange in intact tissue. **(A)** Intravital imaging of ear skin epidermis showing expression of IEbeta-mAmetrine mice in Langerhans cells. Lines in top panel are hair follicles that autofluoresce in all channels. The bottom panel is an enlargement illustrating LC morphology. **(B)** Thymus was isolated from IEbeta-mAmetrine mice and imaged on the two photon microscope ex vivo. The top image is the cortical region near the surface and the lower image is the medullary region deeper in the tissue. **(C)** Cervical lymph nodes were isolated from mice co-expressing CD8beta-LSSSmOrange and IEbeta-mAmetrine and imaged on the two photon microscope ex vivo. mAmetrine expression (green) is localized primarily in peripheral B cell follicles and LSSmOrange (red) expression is localized primarily within the central T cell zone. Two examples from different lymph nodes are shown.

### Expression of CD8beta-LSSmOrange

To determine whether expression of LSSmOrange was a faithful reporter of CD8beta expression expression, we harvested cells from spleen and thymus from WT C57BL/6 mice and from heterozygous (+/-) and homozygous (++) CD8beta-LSSmOrange mice and stained them for endogenous CD8beta. There is good concordance between LSSmOrange and CD8beta in spleen cells and homozygous mice express higher levels of LSSmOrange (Fig 5A,B). In the thymus, LSSmOrange expression is retained in single positive CD4 T cells, presumable due to a longer half-life of LSSmOrange protein compared to CD8 (Fig 5C). This residual LSSmOrange expression is lost in CD4 T cells in the periphery (Fig 5A). To assess the expression of CD8beta-LSSmOrange within intact tissue, we crossed these mice to IEbeta-mAmetrine mice and imaged intact cervical lymph nodes ex vivo on the two-photon microscope. As can be seen in Fig 4C, mAmetrine expression is localized primarily in peripheral follicles containing class II- positive B cells and LSSmOrange expression is localized primarily within the central T cell zone. Taken together, these data indicate that expression of LSSmOrange in CD8beta-LSSmOrange mice accurately reflects expression of endogenous CD8beta expression.

**Figure 5.**
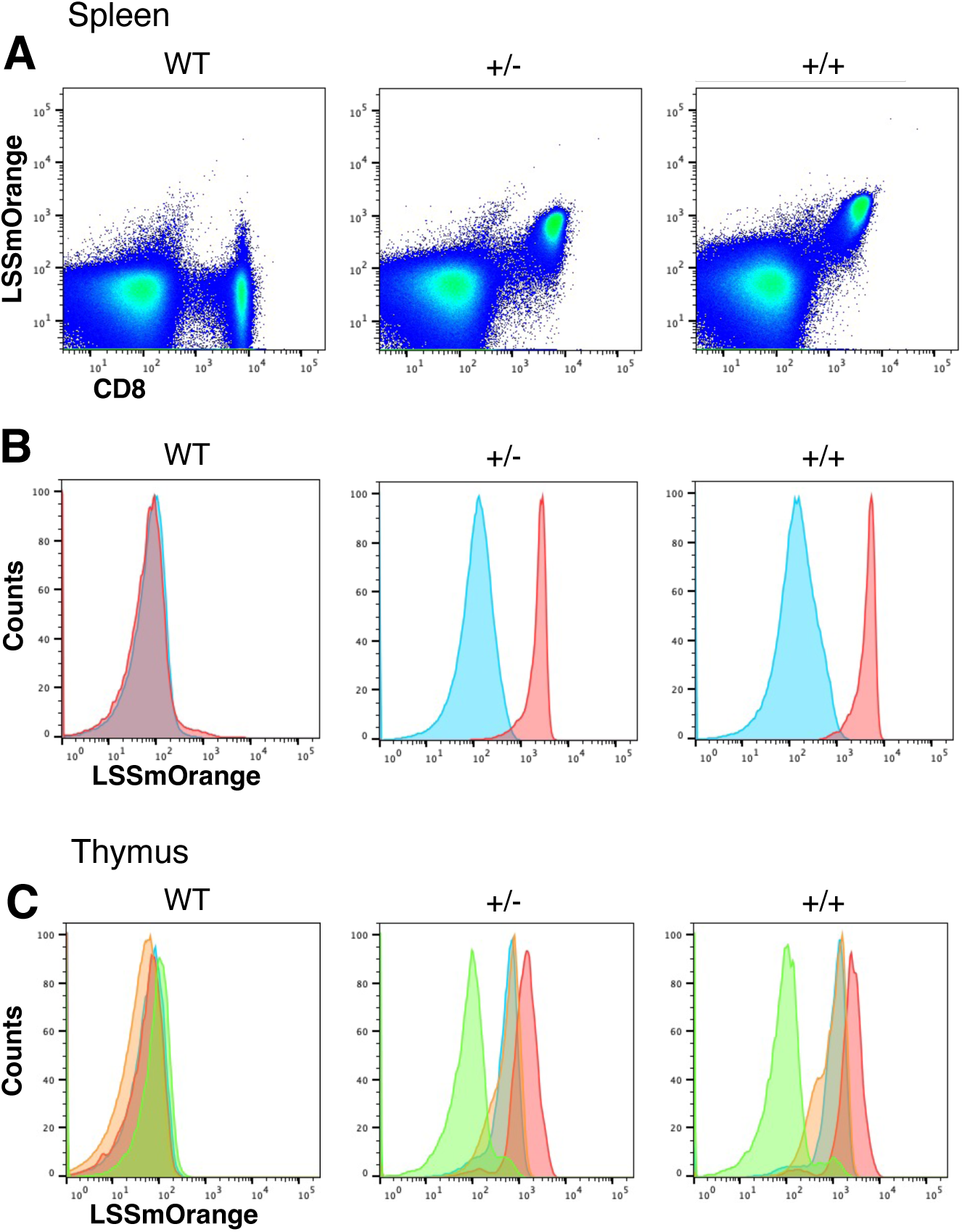
LSSmOrange is a faithful reporter of CD8 expression in CD8beta-LSSmOrange mice. **(A)** Representative flow cytometry of splenocytes isolated from WT mice (left), and from heterozygous (+/-, center) and homozygous (++, right) CD8beta-LSSmOrange mice, stained for CD8beta, and displayed with LSSmOrange expression. **(B)** Histograms of LSSmOrange expression in CD4 (blue) and CD8 (red) cells from spleen of WT, +/-, and ++ mice. **(C)** Histograms of LSSmOrange expression in CD4 single positive (blue), CD8 single positive (red), CD4/CD8 double positive (orange) and CD4/CD8 double negative (green) cells from thymus of WT, +/-, and ++ mice.

Homozygous expression of LSSmOrange did result in a selective decrease in single positive CD8 T cells, starting in the thymus and persisting in the periphery (Fig 6A-C). This decrease in CD8 T cells numbers correlated with a decrease in endogenous CD8beta expression in both singe positive thymocytes and splenic CD8 T cells(Fig 6D-E). This did not appear to be a result of increased expression of the fluorescent protein reporter, as the level of LSSmOrange was proportionally increased in homozygous mice compared to heterozygous mice (Fig 5B, Fig 6F,G). Interestingly, the level of CD8alpha was also increased in homozygous mice (Fig 6 F-G). Conventional CD8 T cells express primarily CD8 alpha-beta heterodimers, rather than CD8 alpha-alpha homodimers, suggesting that beta may be normally expressed in excess to drive primarily heterodimer formation (Devine L et al, 2000, Srinivasan S et al, 2024). Thus, one possible explanation of these observations is that the conformation of the bicistronic CD8- LSSmOrange produced from the knockin construct interferes with translation of CD8, but not LSSmOrange. In heterozygous CD8beta-LSSmOrange mice, there may be little impact on cell surface CD8 expression, as sufficient CD8beta is present to dimerize with CD8alpha. Consistent with this interpretation, there is little impact on CD8 surface expression in hemizygous CD8beta knockout mice (Fung-Leung W-P et al, 1994). However, in CD8beta- LSSmOrange homozygous mice, the level of CD8beta is reduced below this threshold, resulting in a reduction of CD8beta at the cell surface and a slight increase in CD8alpha homodimers. CD8 alpha-beta heterodimers (and not alpha-alpha homodimers) are required for efficient positive selection in the thymus (Cheroutre H & Lambolez F, 2008, Fung-Leung W-P et al, 1994, Gangadharan D & Cheroutre H, 2004), so this loss in CD8beta expression could account for the reduced frequency of CD8 T cells in homozygous CD8beta-LSSmOrange mice.

**Figure 6.**
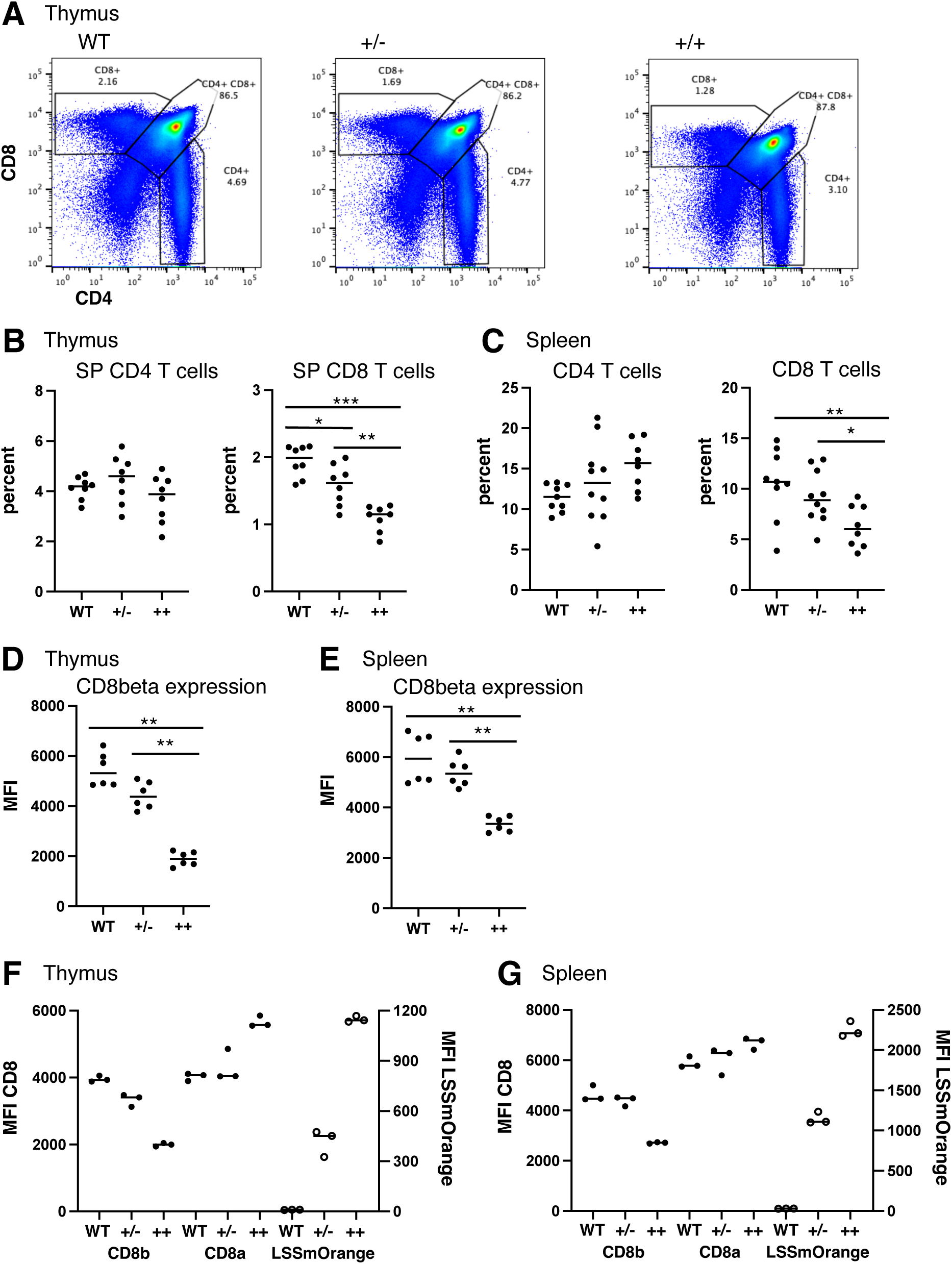
Homozygous expression of CD8beta-LSSmOrange results in loss of CD8 T cells and downregulation of CD8beta expression. **(A)** Representative flow cytometry of thymocytes isolated from WT (left), and from heterozygous (+/-, center) and homozygous (++, right) CD8beta-LSSmOrange mice and stained for CD8 and CD4. Gates and percentages for single positive CD4, single positive CD8 and double positive cells are shown. **(B, C)** Percent of single positive CD4 and CD8 thymocytes (B) or of CD4- and CD8-positive splenocytes (C) from individual WT, +/- and ++ CD8-LSSmOrange mice. **(D, E)** Mean Fluorescence Intensity (MFI) of CD8beta expression in single positive CD8 thymocytes (D) or in CD8-positive splenocytes (E) from individual WT, +/- and ++ CD8-LSSmOrange mice. **(F, G.)** MFI of CD8beta and CD8alpha (closed circles, left hand axis) and MFI or LSSmOrange (open circles, right hand axis) in single positive CD8 thymocytes **(F)** or in CD8-positive splenocytes. **(G)** from individual WT, +/- and ++ CD8-LSSmOrange mice. (***p < 0.0005; **p < 0.005; * P < 0.05).

### The CD8 T cells in CD8beta-LSSmOrange mice are functional

Because LSSmOrange expression can result in a decrease in CD8 cell surface expression and CD8 T cell numbers particularly in homozygous mice, we tested whether the CD8beta- LSSmOrange mice were immunocompetent. WT mice and homozygous and heterozygous CD8-LSSmOrange mice were infected intranasally with influenza virus. After 10 days, the mice were injected with APC-labeled CD45 to stain cells in the vascular (Anderson KG et al, 2014) and lungs were harvested and analyzed by flow cytometry to enumerate the recruitment of CD4 and CD8 cells into the lung tissue (Fig 7A). Lungs from uninfected mice have very few CD4 or CD8 T cells within lung tissue. After infection, similar numbers of both CD4 and CD8 T cells were recruited into the lung in WT, homozygous and heterozygous mice, suggesting that although there are fewer CD8 T cells in the homozygous mice (Fig 6A-C), they are capable of generating a robust response to influenza infection.

**Figure 7.**
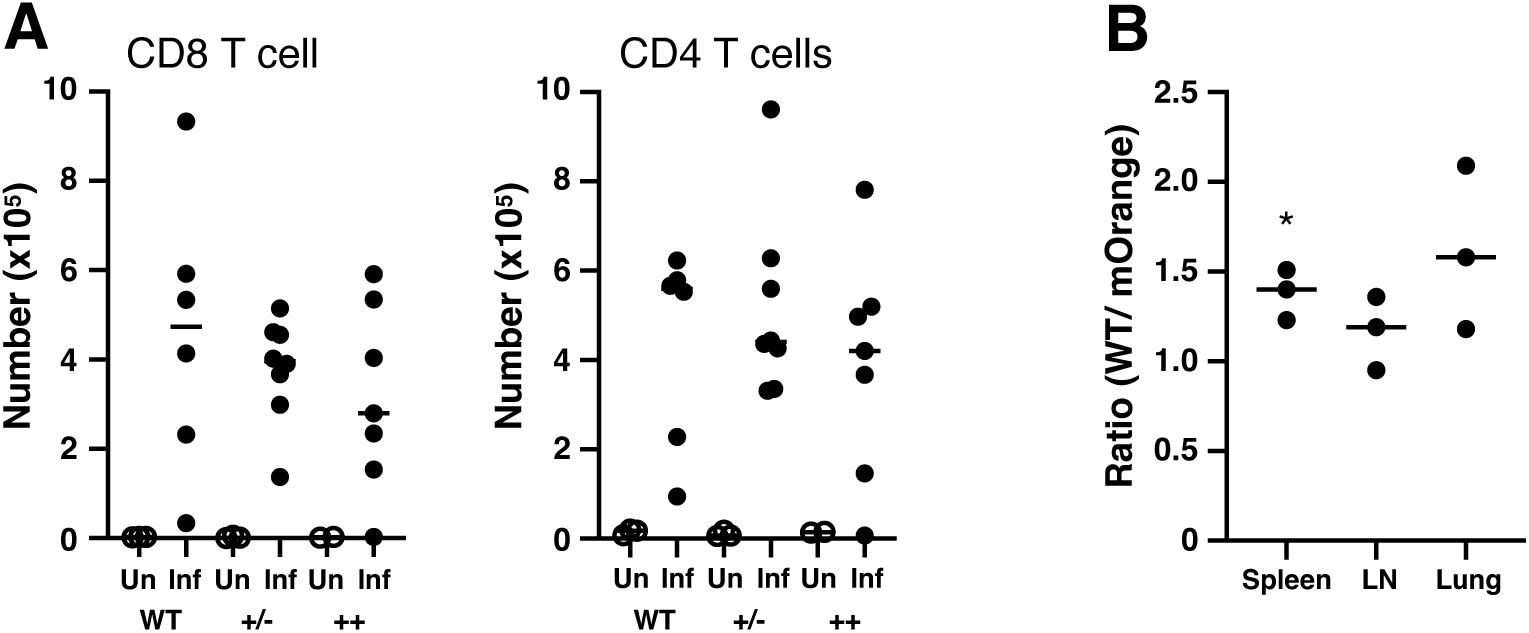
CD8 T cells from CD8beta-LSSmOrange mice can respond to influenza infection. **(A)** WT, heterozygous (+/-), and homozygous CD8beta-LSSmOrange (++) mice were intranasally infected with influenza and on day 10, cells were isolated from lungs from uninfected (Un, open symbols)) and infected (Inf, filled symbols) mice, and the number of lung tissue resident CD8 T cells (left) and CD4 T cells (right) were determined by flow cytometry. (Data combined from 2 separate experiments, n=2-8.There were no significant differences in CD8 or CD4 T cell numbers from WT, +/-, or ++ infected mice). **(B)** T cells were isolated from heterozygous CD8beta-LSSmOrange OTI TCR transgenic and CAG-GFP OTI TCR transgenic mice, co-adoptively transferred into WT mice and the recipient mice were infected with recombinant influenza expressing OVA peptide. The ratio of WT CAG-GFP OTI to CD8-LSSmOrange OTI T cells in spleen, draining lymph nodes, and lung tissue was determined 10 days after infection (3 individual mice, the ratio in spleen, but not lymph node or lung, is significantly different from the starting ratio of 1.02 (* p < 0.05).

To further test the responsiveness of CD8 T cell in CD8beta-LSSmOrange mice, we crossed these mice to OTI TCR transgenic mice. Both WT and CD8beta-LSSmOrange OTI mice showed the same increase in CD8 single positive cells in the thymus cells (supplemental Fig S2A) and increase in numbers of CD8 T cells in the spleen (supplemental Fig S2B) indicative of efficient positive selection of OTI T cells. Furthermore, the majority of the CD8 T cells coexpress Vbeta5 and Valpha2, the TCR chains used by the transgenic OTI TCR (Hogquist KA et al, 1994). To determine whether these OTI T cells were functional, we labeled them with eFlour 450, stimulated them in vitro with antigen and antigen presenting cells, and analyzed the dilution of eFlour 450 as an indicator of cell division. Both WT and CD8beta-LSSmOrange heterozygous OTI cells generated the same pattern of eFluor dilution, indicating that they proliferated equivalently (supplemental Fig S2C). Finally, we co-adoptively transferred naïve WT OTI cells expressing GFP and OTI cells from heterozygous CD8beta-LSSMOrange mice into WT mice and infected the mice with recombinant influenza virus containing the ovalbumin epitope for OTI cells (Reilly EC et al, 2020). At day 10, after vascular staining with anti-CD45, we harvested spleen, draining lymph node, and lung and determined the ratio of GFP-positive WT to CD8beta-LSSmOrange OTI cells within lung tissue (Fig 7B). Although the ratio of WT to CD8beta-LSSmOrange OTI cells is slightly increased, this increase is only statistically significant from the starting ratio in the spleen cells. Collectively, these data indicate that the CD8 T cells in heterozygous CD8beta-LSSmOrange mice are functional.

### Sequential imaging protocol

To follow the kinetics of inflammation with the dermis, we first needed to establish a method to repeatedly image the same region of the mouse ear without inducing additional inflammation. Previous protocols for intravital imaging of the mouse ear utilized hair removal and adhesive tape for immobilization. (Li JL et al, 2012, Li JL et al, 2018, Li S et al, 2018) However, hair removal and repeated tape stripping can disrupt barrier function in the skin and induce inflammation (Nesovic LD et al, 2024). Therefore, we established a procedure to immobilize the ear using only a foam pad and cover slip (see Methods for details). To confirm that repeated imaging using this procedure did not induce inflammation, mice expressing IEbeta-mAmetrine were crossed to mice expressing CD11c-Venus and REX3 (CXCL10-BFP/CXCL9-RFP) and the same area of skin was repeatedly imaged over the course of 26 days. Using vascular and hair follicle landmarks, we are able to localize and collect intravital microscopic images over the same 850μm x 850μm of the ear dermis over the 26 day time course (supplemental Fig S3A).

Class II-positive and CD11c-positive dermal dendritic cells are motile (Kissenpfennig A et al, 2005, Ng LG et al, 2008) and the positions of mAmetrine and Venus expressing cells do change over time. However, there is no apparent induction of CXCL10 expression (BFP, blue) or increase in the number of MHC class II-positive cells (IEbeta-Ametrine, green) or CD11c- positive cells (yellow), indicative of minimal tissue response to the repeated imaging protocol (supplemental Fig S3B-C).

### Inflammation induced activation of Langerhans cells

As a first test to utilize the IEbeta-mAmetrine mice to monitor cellular changes following induction of an inflammatory response, we imaged the epidermal, LC layer of the ear skin prior to and two days after induction of inflammation with CFA. Once again, we were able to localize the same tissue area at each time point (Fig 8A). CFA induced inflammation resulted in a reduction in the density (Fig 8B) and number (Fig 8D) of LC within the skin. In addition, there was a noticeable change in LC morphology, with a retraction of dendrites and rounding of the cell body (Fig 8C). These data are consistent with LC activation response to innate signals in CFA resulting in changes in motility and ultimate migration from the skin to the draining lymph node. (Eidsmo L et al, 2009, Kissenpfennig A et al, 2005)

**Figure 8.**
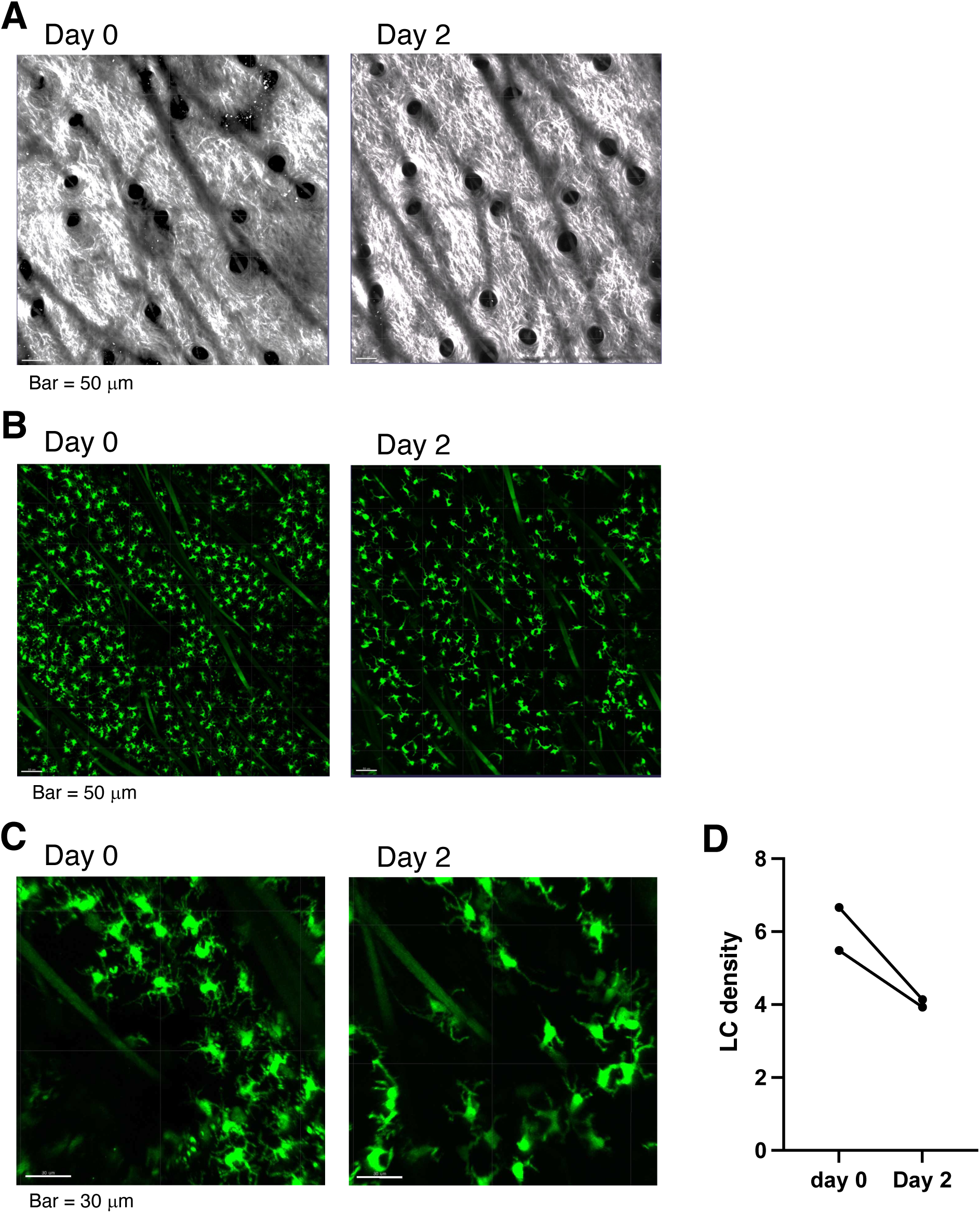
Activation of Langerhans cells after intradermal immunization with OVA/CFA. IEbeta-mAmetrine mice was immunized in the dermis of the ear with OVA/CFA containing a tracer of Alexa647-OVA. Four hours later, the OVA/CFA depot was localized, an area proximal to the depot and distal to the site if injection was chosen, and an image in the epidermal layer was collected intravitally on the two photon microscope. Two days later the same area was identified and re-imaged. **(A)** Relative position of hair follicles (dark circles) in the second harmonic generation image confirms imaging of the same location. **(B)** Fluorescent image of IEbeta-mAmetrine expression in Langerhans cells. Note the reduced number of LC on day 2. The green lines are autofluorescent hair follicles. **(C)** Enlarged image of LC. Note the change in morphology (retraction of dendrites and enlargement of cell body) of LC on day 2. **(D)** Quantitation of the density of LC (# LC/mm^2^) from two separate experiments.

### Induction and resolution of peripheral activation clusters

To follow the induction of inflammation in the skin further, we crossed the IEb-mAmetrine mice to mice expressing CD11c-Venus and REX3 expressing CXCL10-BFP/CXCL9-RFP. The mice were intradermally immunized with OVA in CFA and then using vascular and hair follicle landmarks the same region of the skin was sequentially imaged from 6 to 33 days (Fig 9A). As has been previously reported, CFA immunization induces clusters of CXCL10 expressing cells, termed T cell activation niches (Bala N et al, 2022, Prizant H et al, 2021). During this time course, we were able to observe the expansion and contraction of the chemokine-expressing cell clusters that peak on day 12 (Figs 9B and 10A). In addition, based on differential expression of IEbeta-mAmetrine, CD11c-YFP, and CXCL10-BFP, we were able to identify 4 different populations of class II expressing cells (Figs 9C and 10B-E). There are few class II-positive cells in the tissue on day 6, when the CXCL10+ cluster have already begun to form. All 4 class II positive populations increase in number, localizing primarily within the clusters and peak on day 12. However, the two CXCL10-positive populations predominate during these early time points (Fig 10C). From day 12 to day 33, the size of the clusters diminish, suggesting an abatement of the inflammatory response (Figs 9B and 10A). In contrast, the total number of class II positives cells continue to increase (Fig 10B). By day 33, the majority of class II positive cells reside outside of the clusters. However, cells expressing all three markers (class II, CD11c, and CXCL10) remain primarily within the clusters (Fig 10D).

**Figure 9.**
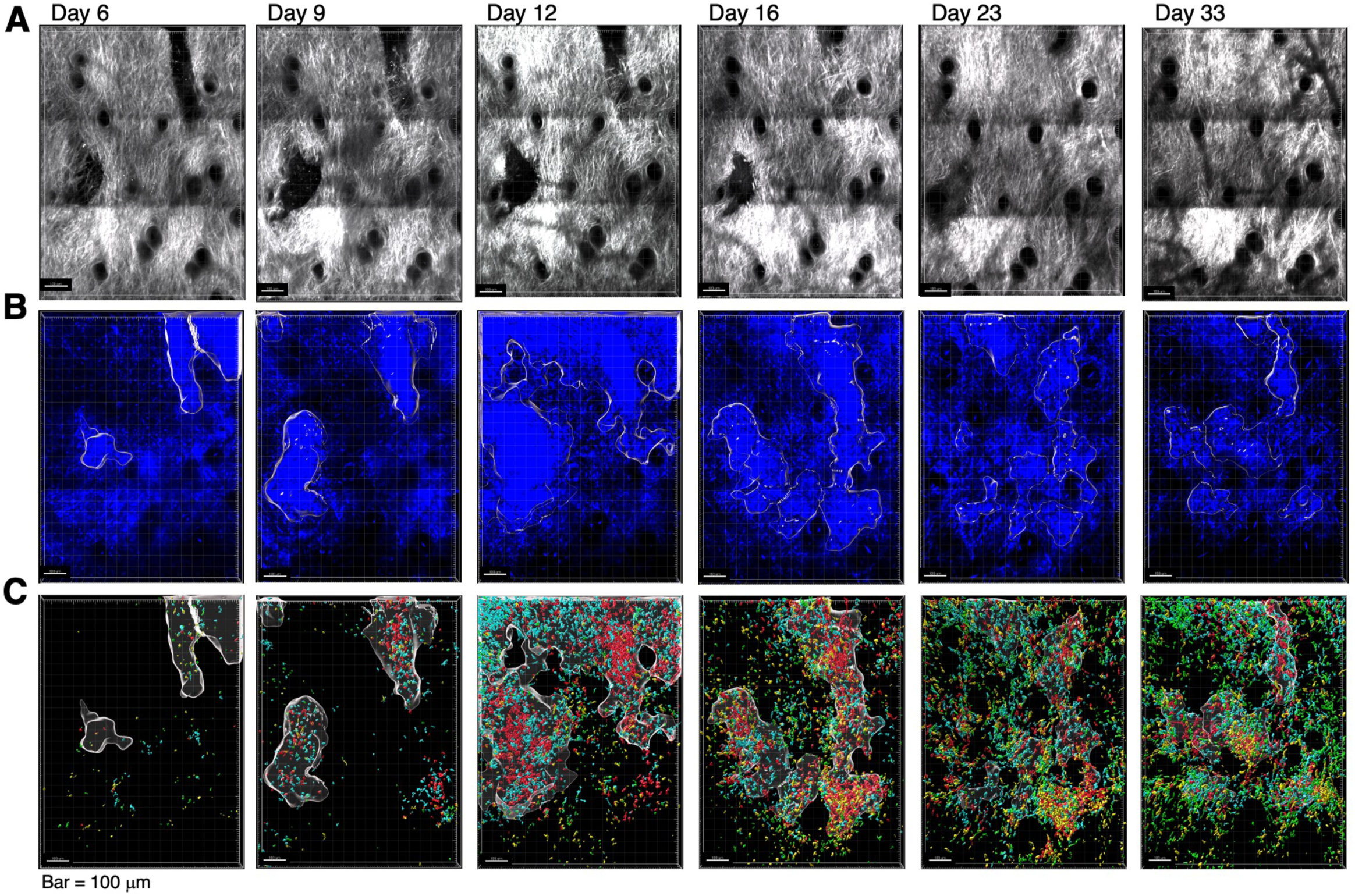
Sequential intravital imaging of class II expressing cells within activation clusters in inflamed ear dermis. IEbeta-mAmetrine mice were crossed to CD11c-YFP and REX3 mice and intradermally immunized with ovalbumin in CFA in the ear pinna and the same area of tissue was sequentially imaged by two-photon microscopy on days 6, 9, 12, 16, 23, and 33. **(A)** Relative position of hair follicles (dark circles) in the second harmonic generation image confirms imaging of the same location. **(B)** Clusters of CXCL10-BFP expressing cells were identified by nearest neighbor analysis (20 cells within radius of 50mm) and highlighted with white border. **(C)** Different populations of MHC class II positive cells were based on differential expression of fluorescent proteins. Green, class II+ only; Yellow, class II+ and CD11c+; Blue, class II+ and CXCL10+; red, class II, CXCL10+ and CD11c+; white outline, clusters of CXCL10+ cells. Representative data from two independent experiments.

**Figure 10.**
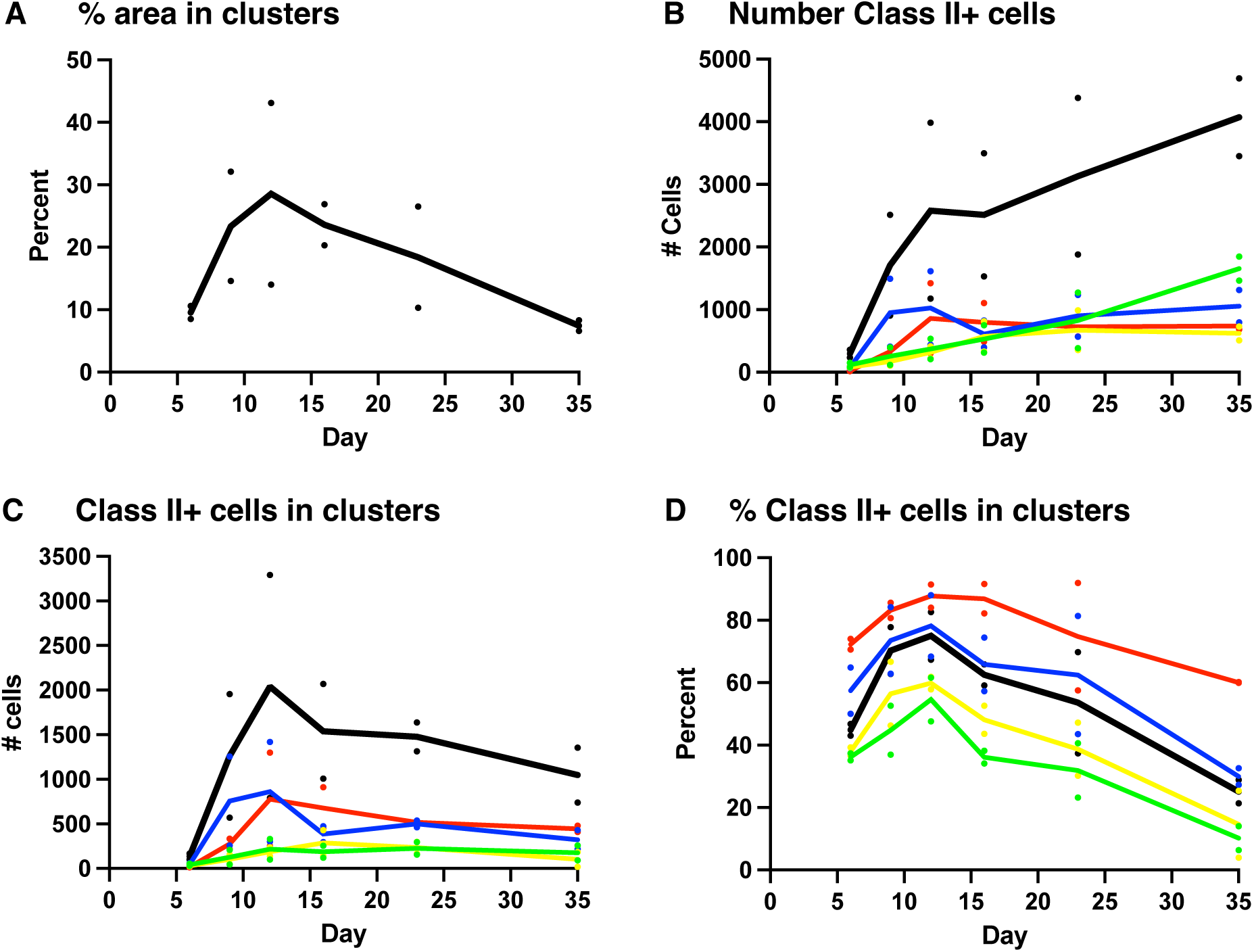
Enumeration of class II-positive cells within clusters of CXCL10-expressing cells. CXCL10 clusters and class II positive cells were identified from the images shown in Fig 9 and the class II+ cells expressing different combinations of CD11c and CXCL10 within and outside the CXCL10 clusters was determined. **(A)** Percent of area contained within CXCL10 clusters. **(B)** Number of class II+ cells with entire image. **(C)** Number of class II+ cells with CXCL10 clusters. **(D)** Percent of class II+ cells within clusters. **(B-D)** Total number of class II+ cells (black); class II+ only (green); class II+ and CD11c+ (yellow); class II+ and CXCL10+ (blue) and class II, CXCL10+ and CD11c+ (red). Data are average of two separate experiments with individual data points for each time point shown.

### Kinetics of T cell localization within peripheral activation clusters

To image recruitment of T cells, we adoptively transferred OTII Th1 cells expressing OFP into IEb-mAmetrine/CD11c-Venus/REX3 mice and immunized them with OVA/CFA. As above, we imagined the same area of the skin during the sequential time points (Fig 11A) and observed the formation, expansion, and contraction of the of the chemokine-expressing cell clusters (Fig 11B and 12A) and the recruitment of different MHC class II-positive cells into the tissue and localization within the CXCL10 clusters (Fig 11C and 12B-C). T cells increase dramatically within the tissue from day 3 to day 6 and then slowly wane, with few T cells remaining by about day 30 (Fig 12D). The percentage of T cells within the CXCL10 clusters peaks a bit later at about 60% on day 12 (Fig 12E). Overall, the kinetics of cluster formation and resolution, the recruitment of class II-positive cells into the tissue, and the accumulation of MHC class II positive cells into and dispersal from CXCL10 cluster is similar in the presence and absence of exogenous T cells (Fig 13), suggesting that fluorescently labeled T cells are a reasonable representation of the endogenous T cell response. Although, the addition of the pre-activated Th1 cells does enhances the early expression of class II positive cells within the clusters, possible due to the increased availability of IFN-gamma (Fig 13B). Notably, the presumptive monocyte-dendritic cells, that co-express MHC class II, CD11c, and CXCL10, selectively persist within the clusters in the presence and absence of exogenous T cells (red lines in Fig 13C), even though at this point most of the overall number of T cells is reduced and the remaining T cells are no longer enriched within the clusters (Fig 12 D-E).

**Figure 11.**
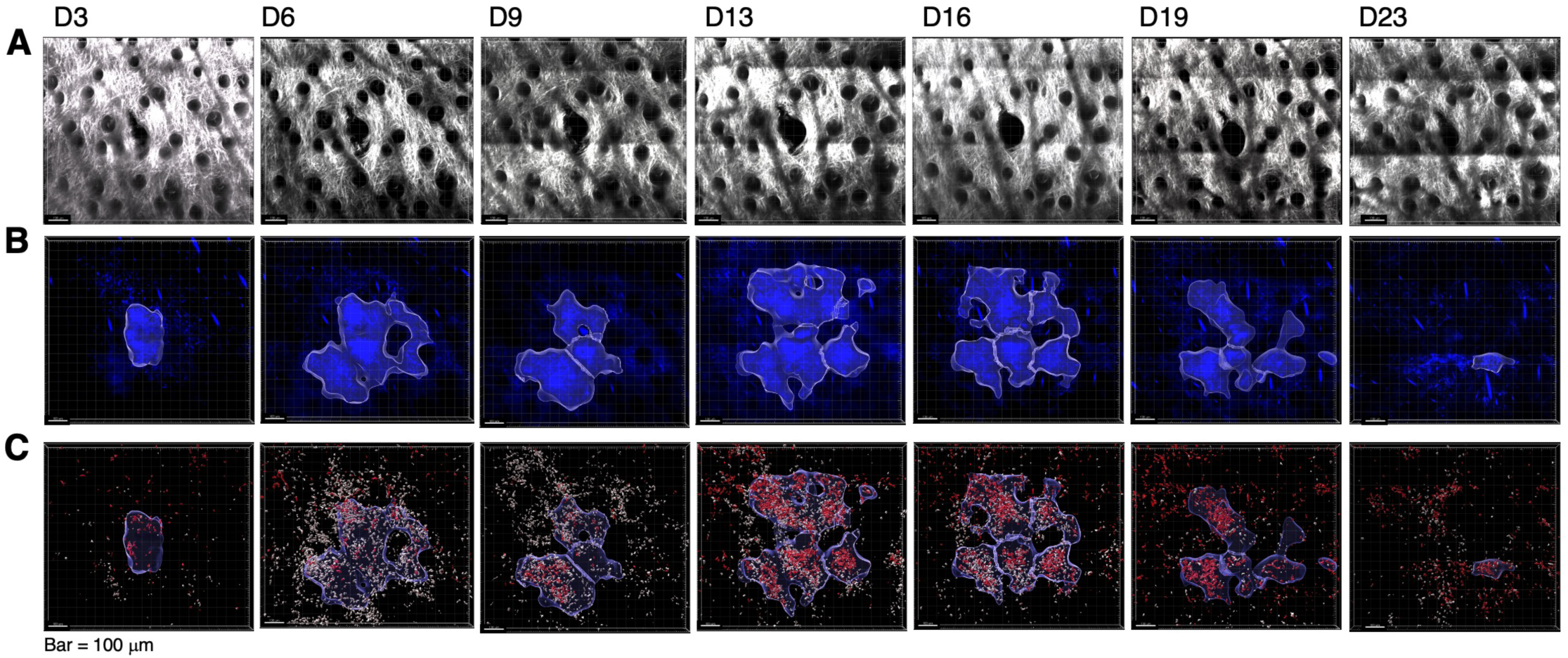
Sequential intravital imaging of T cell recruitment to activation clusters in inflamed ear dermis. IEbeta-mAmetrine mice were crossed to CD11c-YFP and REX3 mice and were adoptively transferred with 1x10^6^ in vitro activated OTII Th1 cells expressing OFP. The mice were intradermally immunized with ovalbumin in CFA in the ear pinna and the same area of tissue was sequentially imaged by two-photon microscopy on days 3, 6, 9, 13, 16, 19, and 23. **(A)** Relative position of hair follicles (dark circles) in the second harmonic generation image confirms imaging of the same location. **(B)** Clusters of CXCL10-BFP expressing cells were identified by nearest neighbor analysis (20 cells within radius of 50mm) and highlighted with white border. **(C)** All class II positive cells are shown in white, OTII Th1 cells are shown in red, and the outline of the cluster defined in **B** is shown. Representative data from one of three experiments. Data from the other two experiments is show in supplemental Figure S4.

**Figure 12.**
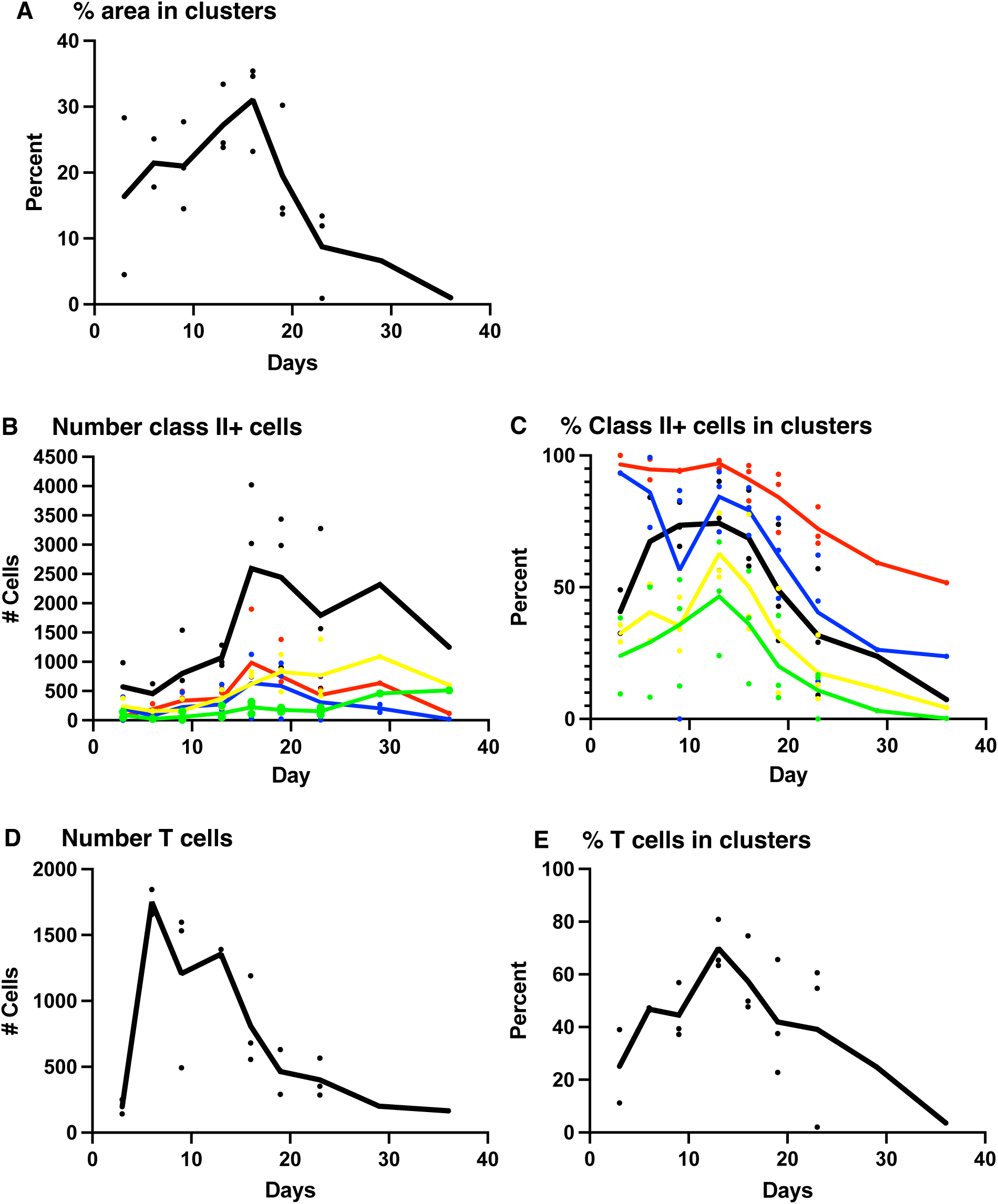
Enumeration of class II-positive cells and Th1 T cells within clusters of CXCL10-expressing cells. CXCL10 cluster, Th1 T cells, and class II positive cells and were identified from the images shown in Fig 11 and supplemental Fig S4. **(A)** Percent of area contained within CXCL10 clusters. **(B)** Number of class II+ cells with entire image **(C)** Percent of class II+ cells within clusters. **(D)** Number of Th1 T cells with entire image. **(E)** Percent of T cells within clusters. **(B-C)** Total number of class II+ cells (black); class II+ only (green); class II+ and CD11c+ (yellow); class II+ and CXCL10+ (blue) and class II, CXCL10+ and CD11c+ (red). Data are mean of three separate experiments with individual data points for each time point shown. Note there are only 2 data points for day 3 and day 6 and single data point for days 29 and 36.

**Figure 13.**
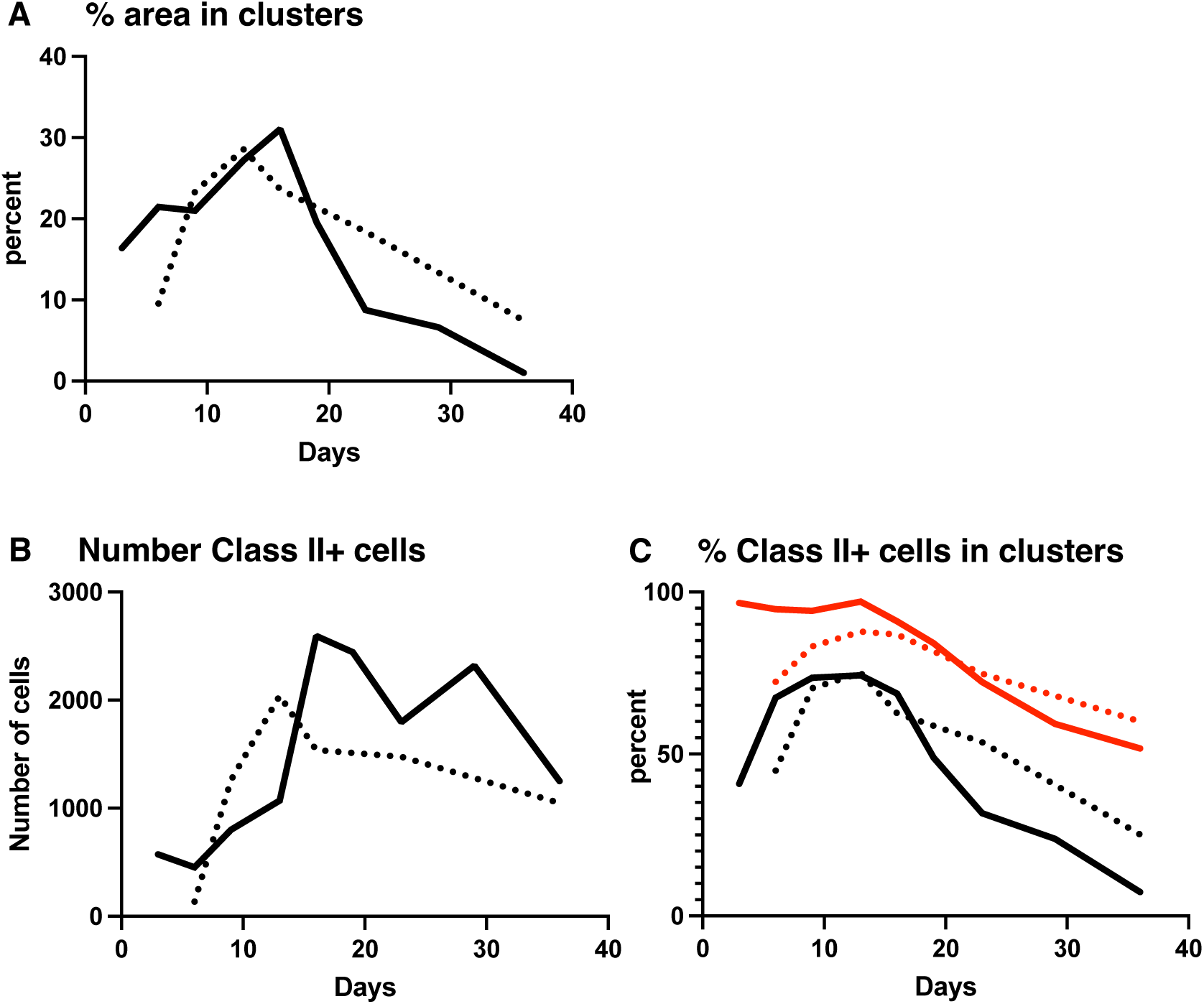
Comparison of cluster formation in the presence and absence of exogenous T cells. Data from Fig 10 (without exogenous T cells, dotted lines) and Fig 12 (with exogenous T cells, solid lines) are overlaid. **(A)** Percent of area contained within CXCL10 clusters. **(B)** Number of class II+ cells with entire image. **(C)** Percent of total class II+ cells within clusters (lack lines) and percent of class II+, CD11c+, CXCL10+ triple positive cells (red lines).

In sum, we have developed two new fluorescent protein mouse models that faithfully reflect the expression of MHC class II (IEbeta-mAmetrine) and conventional CD8-positive T cells (CD8beta-LSSmOrange) and have submitted these strains to Jackson labs for distrubition to the community. We have also established a method for repeated imaging of the same region of the dermis and have used the IEbeta-Ametrine mice crossed to CD11c-Venus and REX3 (CLCL10- BFP) mice and adoptively transferred with OFP labeled to cells to monitor the initiation, amplification, and resolution of an antigen-specific inflammatory response within skin tissue.

Utilization of a dual laser two photon microscope and spectral unmixing enabled us to simultaneously identify and enumerate 5 different populations of cells expressing different combinations of fluorescent proteins.

## Methods

### Mice

To generate IEbeta-Ametrine mice, a cassette containing the mAmetrine coding sequence fused to a rabbit beta globin 3’UTR/poly A addition sequence (Fig 1 and supplemental Table S2), a sgRNA [5′-TCCTCTCCTGCAGCATGGTG (TGG)-3′], and purified CAS9 were microinjected into fertilized eggs isolated from inbred C57BL/6 mice to knock Ametrine into the translational start site of the endogenous class II IEbeta gene. Offspring were screened by PCR and accurate knockin was confirmed by DNA sequencing. The mice were backcrossed to C57BL/6-albino mice (Jackson Laboratory, Bar Harbor, ME) and maintained in a specific-pathogen-free facility at the University of Rochester Medical Center according to institutional guidelines. These mice are now available from The Jackson Laboratory (Stock No. 039963 IEbeta-mAmetrine)

To generate CD8-LSSmOrange mice, a cassette containing an IRES fused to the LSSmOrange coding sequence (Fig 1 and supplemental Table S3), a sgRNA [5′- CTAGCAGGCTATCAGTGTTG (TGG)-3′], and purified CAS9 were microinjected into fertilized eggs isolated from inbred C57BL/6 mice to knock IRES-LSSmOrange into the 3’UTR four nucleotides downstream from the termination codon of the endogenous CD8beta gene.

Offspring were screened by PCR and accurate knockin was confirmed by DNA sequencing. The mice were backcrossed to C57BL/6-albino mice (Jackson Laboratory) and maintained in a specific-pathogen-free facility at the University of Rochester Medical Center according to institutional guidelines. These mice are now available from The Jackson Laboratory (Stock No. 039964 CD8beta-LSSmOrange)

### Tissue preparations and flow cytometry

Single cell suspensions were prepared from thymus, spleen, and lymph nodes by passing through a 40 μm nylon mesh filter. Lung cells suspensions were prepared by tissue mincing, enzymatic digestion in 1 mg/ml Type II collagenase (Gibco) and 30 μg/ml DNase I (Roche) for 45 minutes at 37 °C and then by passing through a 40 μm nylon mesh filter. Single cell suspensions were treated with ACK lysis buffer (0.15M NH4Cl/1mM KHCO3/0.1mMNa2-EDTA, pH 7.2) for 5 min to deplete red blood cells. Staining, washes, and resuspensions were performed in 1X PBS with 5% newborn calf serum (Hyclone).

Cells were pre-stained with Ghost Dye Violet 510 or Ghost dye Blue 516 (Tonbo Biosciences) to exclude dead cells and with anti-CD16/CD32 (2.4G2, Tonbo) to block FcR binding. Cells were then stained with fluorescently-labeled antibodies as indicated in figure legends. Cells were run on an LSR-Fortessa or Symphony A1 flow cytometer and data were analyzed using FlowJo software.

### T cell assays

CD8beta-LSSmOrange mice were crossed to OTI TCR transgenic mice and thymus and spleen were stained with antibodies to CD8, TCR Valpha2, and TCR Vbeta5 to confirm that LSSmOrange-positive CD8 T cells expressing the OTI TCR were appropriately selected. T cells were isolated from spleens and lymph nodes harvested from WT and from heterozygous CD8beta-LSSmOrange mice, were labeled with eFlour 450 (eBioscience) and were stimulated in vitro with 2μg/ml OVA peptide and irradiated spleen cells. On subsequent days, T cell division was monitored by dilution of eFlour 450 fluorescence intensity on the flow cytometer.

For influenza T cell responses, mice were anesthetized with 2,2,2-tribromoethanol (Avertin) via intraperitoneal injection and infected intranasally with 225,000-400,000 egg infectious doses (EID_50_) of influenza A/New Caledonia/20/1999 (H1N1) virus. After 10 days, animals were anesthetized with isoflurane and injected retro-orbitally with 3 μg APC-labelled anti-mouse CD45 (30-F11; Tonbo) and sacrificed 3 minutes later to selectively label vascular cells (Anderson KG et al, 2014). Cells were isolated from lungs of infected and uninfected mice and the number of interstitial CD8 and CD4 T cells were enumerated by flow cytometry.

For the competitive adoptive transfer experiment, spleen and lymph node cells were isolated from heterozygous CD8-LSSmOrange or CAG-GFP (Jackson Laboratory) mice co-expressing the OTI TCR transgene. The cells were mixed to yield equal numbers of WT GFP and LSSmOrange CD8 OTI cells and 1.1x10^6^ total cells containing 2x10^5^ total OTI cells were adoptively transferred by i.p. injection into WT C57BL/6 mice. The mice were then infected with 3x10^3^ EID_50_ HKx31-OVAI expressing the ovalbumin (OVA^257-264^ SIINFEKL) peptide in the stalk of neuraminidase (Reilly EC et al, 2020) and the ratio of CAG-GFP OTI to CD8-LSSm-Orange OTI T cells in spleen, draining lymph nodes, and lungs was determined 10 days after infection.

### Two Photon microscopic imaging

Images were acquired with an Olympus FV1000-AOM multiphoton system equipped for 4-color detection. Fluorescence was collected with an Olympus XLPlanN 25x objective (numerical aperture, 1.05) and was detected with three proprietary photomultipliers. Fluorescence excitation was achieved by sequential scans with a Spectra-Physics DeepSee-MaiTai HP Ti:Sapphire laser and a Spectra-Physics InsightX3 laser. Laser, dichroic mirror, and bandpass filter configurations as show in supplemental Table S1.

For imaging intact tissue, thymus was isolated from IEbeta-mAmetrine mice and cervical lymph nodes were isolated from mice co-expressing IEbeta-mAmetrine and CD8beta-LSSmOrange. The tissue was mounted on slides, a z series was collected through the tissue and single z planes were chosen for display.

For imaging epidermal and dermal layers of mouse ears, mice were anesthetized with isoflurane (induction 4%; maintenance 1.5%, in room air) with an isoflurane vaporizer-ventilation machine (M3000R; Lei Medical). Once the mice were anesthetized, the ear pinna was immobilized between a polyester/acrylic resin foam pad (Home Techpro Vacuum Tech rug gripper) and a cover slip without using adhesive or hair removal to avoid any barrier disruption. Body temperature was maintained with a heated water pad (Kent Scientific) and a heating block (WPI). The microscope objective was heated (Bioptechs) to 40°C to maintain constant dermal temperature during imaging.

### Intravital multiphoton imaging

IEbeta-mAmetrine mice were crossed to CD11c-Venus (Lindquist RL et al, 2004) (Jackson Lab) mice and to REX3 (CXCL10-BFP/CXCL9-RFP) mice (Groom JR et al, 2012) and the triple positive mice were maintained on a C57/BL6-albino background. The mice were immunized by intradermal injection into ear pinna of 1.76 ug ovalbumin emulsified in CFA (Complete Freund’s Adjuvant). In some experiments, Alexa 647-ovalbumin (Thermo Fisher) was added to label the antigen depot. For adoptive transfer experiments, OTII TCR transgenic mice were crossed to CAG-mOFP mice (Gossa S et al, 2014) and 1-4x10^6^ CD4 Th1 cells were adoptive transferred into IEbeta-mAmetrine/CD11c-YFP/REX3 mice prior to immunization. For in vitro generation of effector Th1 cells, T cells were harvested from lymph nodes and spleens of OT-II TCR transgenic mice, and stimulated with 2 μg/ml ovalbumin peptide and irradiated spleen cells in the presence of IL-2 (10 units/ml, NCI BRB Preclinical Biologics Repository) IL-12 (20ng/ml; Peprotech), and anti-IL-4 (40 μg/ml; 11B11, NCI BRB Preclinical Biologics Repository).

For sequential imaging of the same tissue area, the initial localization is based on the organization of the vasculature within a full ear image and then by sequential reiteration of vascular and hair follicle orientation, the precise location can be established for subsequent imaging. At each time point, an 1100μm x 1100μm or an 850μm x 850μm region was stitched together from 3x3 or 4x4 tiles using Olympus software. Relying on the distribution of collagen fibers detected by second harmonic images and position of hair follicles, images were cropped along the x, y and z axes to maximize overlap between time points. Clusters of CXCL10- expressing cells (BFP-positive) were defined in 3D using a Python code that utilizes the unsupervised machine learning algorithm DBSCAN (Density-based spatial clustering of applications with noise) (Ester M et al, 1996). There are two parameters to the algorithm; maximum distance between points, radius (R) and minimum number of neighbors (N) within R to define as part of the cluster. Density cutoffs of R = 50μm and N = 20 neighbors were applied to define the CXCL10 cluster. The images were then spectrally unmixed using Lumos software (McRae TD et al, 2019) to eliminate bleed over between mAmetrine and BFP or between OFP, Venus, BFP, and RFP and channels representing MHC class II only (mAmetrine only); class II and CD11c (mAmetrine and Venus), class II and CXCL10 (mAmetrine and BFP); class II and CD11c and CXCL10 (mAmetrine, Venus, and BFP) and, when present T cells (mOFP only) were separated. Surfaces were generated in 3D software from Imaris (Bitplane) to identify individual cells in each channel and the number of each cell population within and without the CXCL10 clusters was determined. Statistical tests were done using Mann-Whitney in GraphPad Prism.

## Acknowledgements

We thank Drs. Andrea Sant and David Topham for sharing Influenza viruse; Drs. Dorian McGavern and Dr. Andrew Luster for providing CAG-OFP and REX3 mice, respectively; Dr Lin Gan for generating the knockin mice; and Dr. Alex Wells for Langerhans Cell discussions. We acknowledge the support of the URMC flow cytometry advanced light microscopy cores. This work was supported by National Institutes of Health (NIAID) P01 AI02851.

## Author contributions

J Miller: conceptualization, formal analysis, funding acquisition, investigation, methodology, project administration, supervision, validation, visualization, writing – original draft, review & editing D Oleksyn: conceptualization, formal analysis, investigation, methodology, software, validation, visualization, writing – review & editing

The authors have no conflict of interests.

---

**Supplemental Figure S1.**
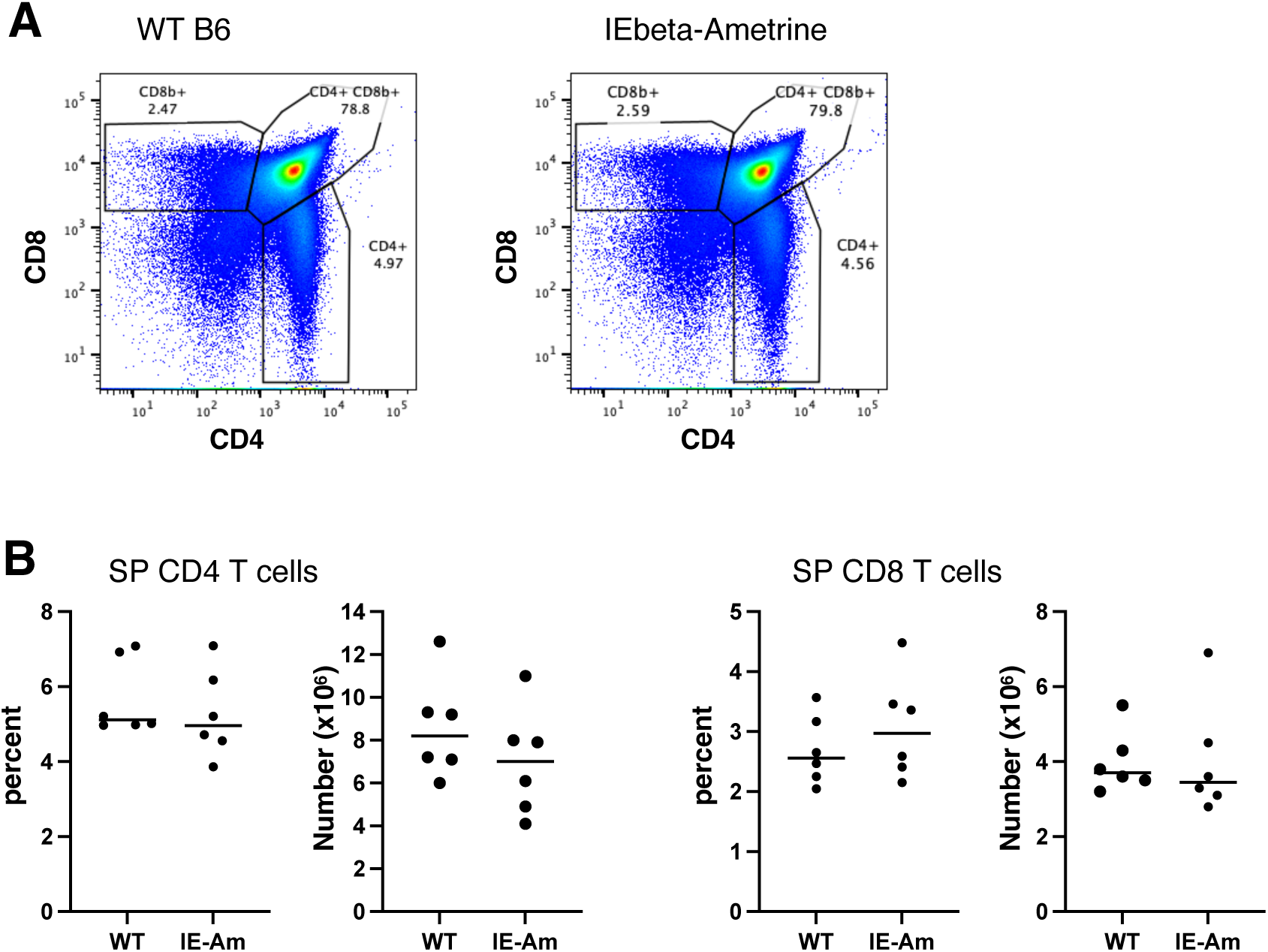
mAmetrine expression in IEbeta-mAmetrine mice does not alter the generation of single positive CD4 and CD8 T cells in the thymus. **(A)** Representative flow cytometry of thymocytes isolated from WT mice (left) and IEbeta-mAmetrine mice (right) stained for CD8 and CD4. Gates and percentages for single positive CD4, single positive CD8 and double positive cells are shown. **(B)** Percent and number of single positive CD4 and CD8 thymocytes from individual WT and IEbeta-Ametrine (IE-Am) mice (data are from 3 mice each from 2 separate experiments, no significant differences).

**Supplemental Figure S2.**
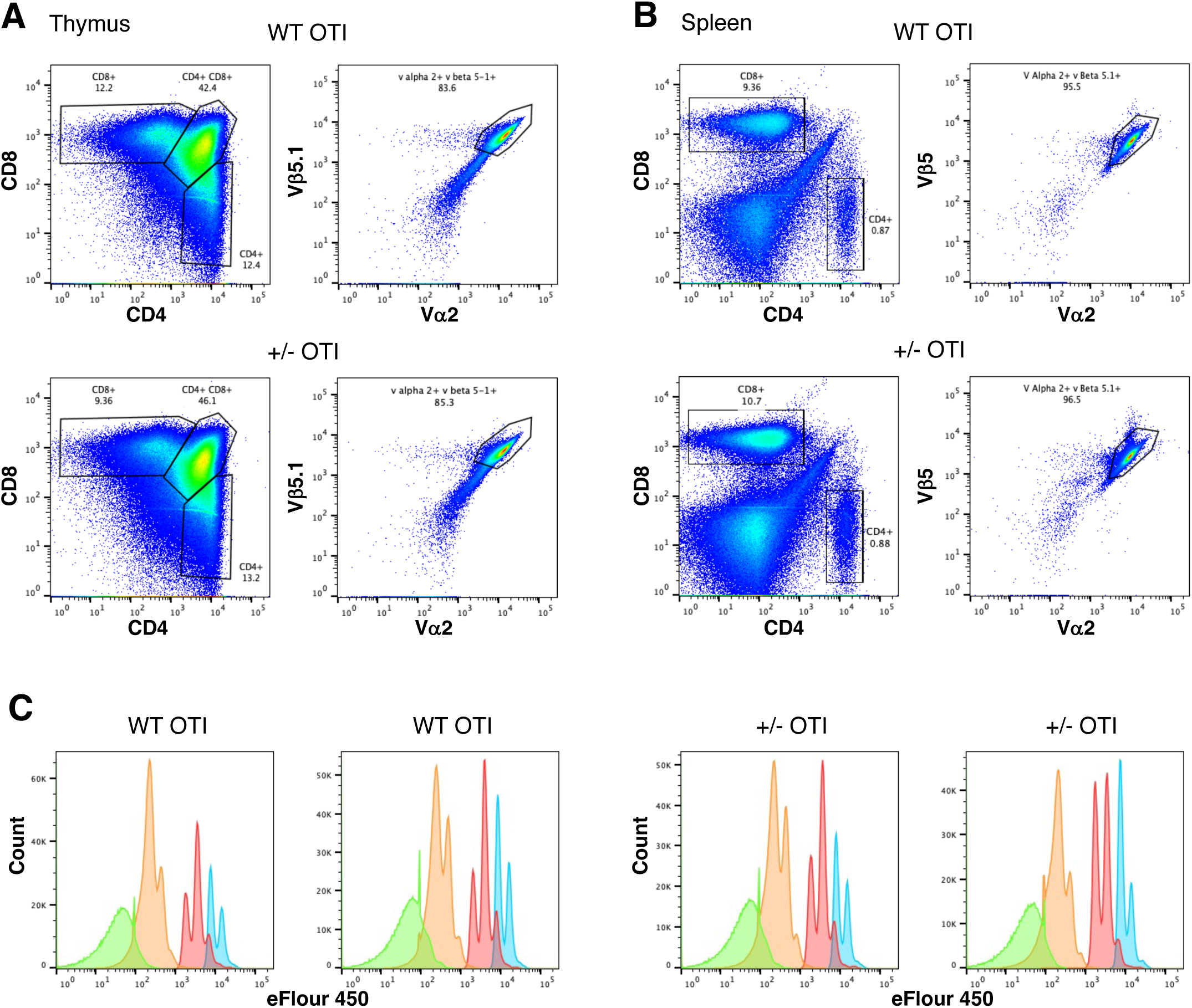
Expression of CD8beta-LSSmOrange does not interfere with thymic development of OTI T cells nor with the ability of OTI T cells to proliferate to antigen in vitro. **(A,B)** Thymus **(A)** and spleen **(B)** were harvested from WT OTI mice and from heterozygous CD8beta-LSSMOrange (+/-) OTI and expression of CD4 and CD8 is shown on the left and co-expression of Vbeta5 and Valpha2 in single positive CD8 T cells is shown on the right. **(C)** CD8 T cells from WT OTI and heterozygous CD8beta-LSSmOrange OTI (+/-) mice were loaded with eFlour 450 dye and stimulated in vitro with irradiated spleen cells and OVA peptide. Dilution of eFlour 450 was determined at days 2 (blue), 3 (red), 4 (orange) and 7 (green). Cultures from 2 independent WT and +/- mice are shown.

**Supplemental Figure S3.**
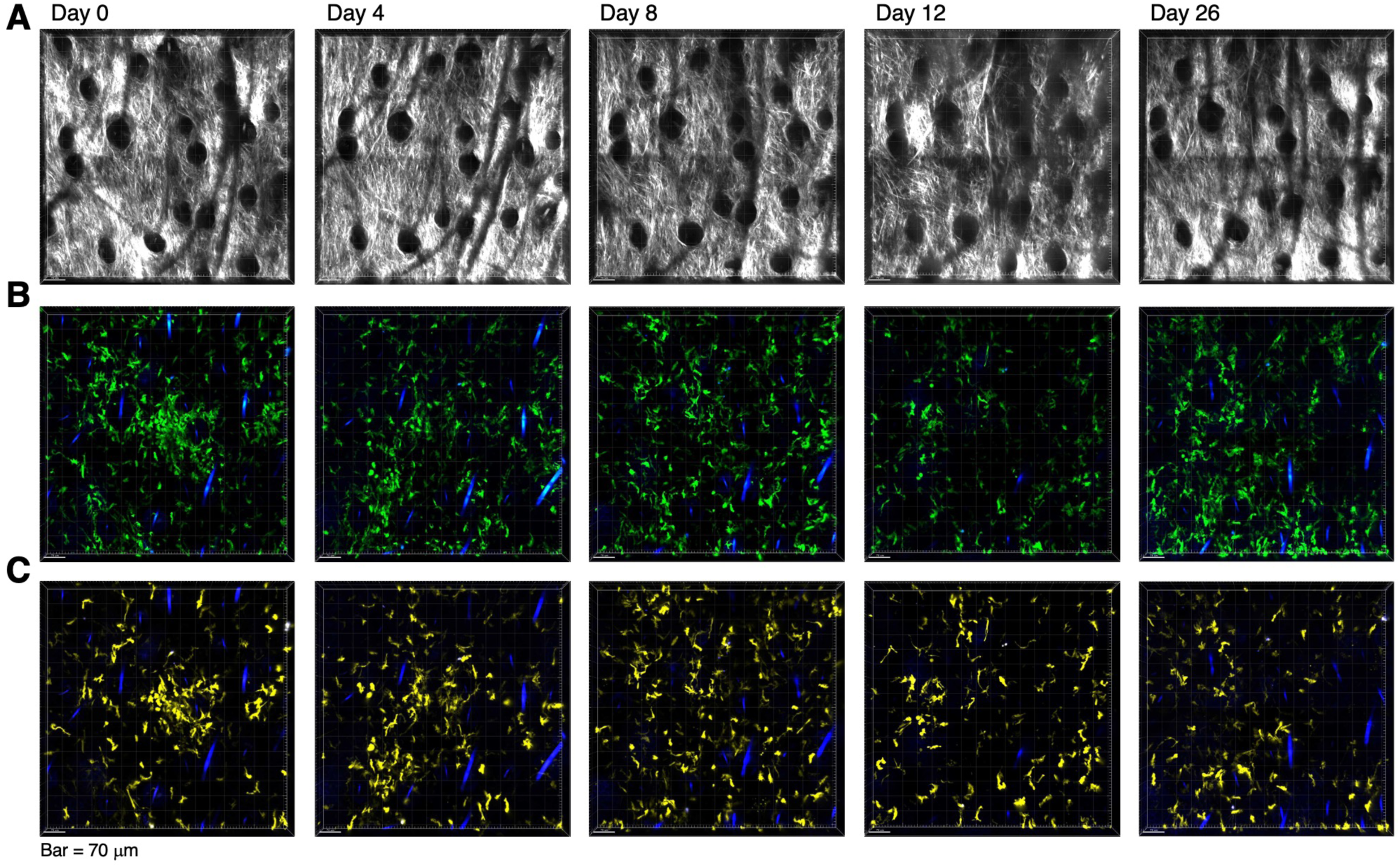
Sequential imaging of ear dermis does not induce inflammation. IEbeta-mAmetrine mice were crossed to CD11c-Venus and REX3 (CXCL10-BFP/CXCL9-RFP) mice. The same site in the ear dermis of unimmunized mice was intravitally imaged on days 0, 4, 8, 12, and 26. **(A)** Relative position of hair follicles (dark circles) in the second harmonic generation image confirms imaging of the same location. **(B)** mAmetrine (green) and BFP (blue) image. **(C)** Venus (yellow) and BFP (blue) image. The blue lines in **B** and **C** are hair follicles that autofluoresce in multiple channels. Representative data from one of 4 separate regions from 2 different mice.

**Supplemental Figure S4.**
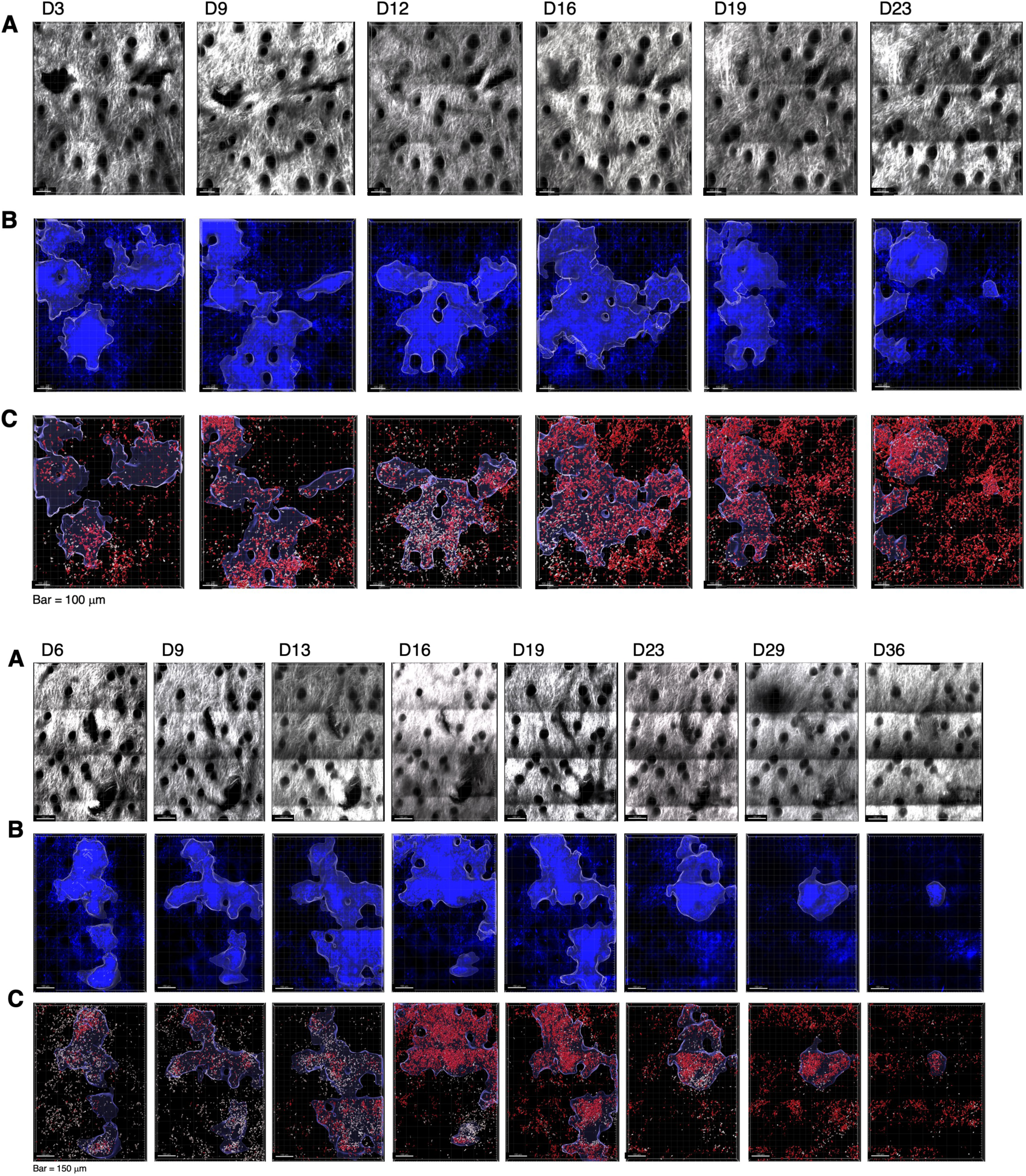
Sequential intravital imaging of T cell recruitment to activation clusters in inflamed ear dermis. Replicate experiments of Fig 11 and quantified in Fig 12 using 4x10^6^ OTII Th1 cells and imaged on days 3, 9, 12, 16, 19, and 23 (top) and 1x10^6^ OTII Th1 and imaged on days 6, 9, 13, 16, 19, 23,29, and 36 (bottom). **(A)** Second harmonic generation images. **(B)** Clusters of CXCL10-BFP expressing cells highlighted with white border. **(C)** All class II positive cells in white, OTII Th1 cells in red, and the outline of the CXCL3-BFP cluster are shown.

**Supplemental Table 1.**
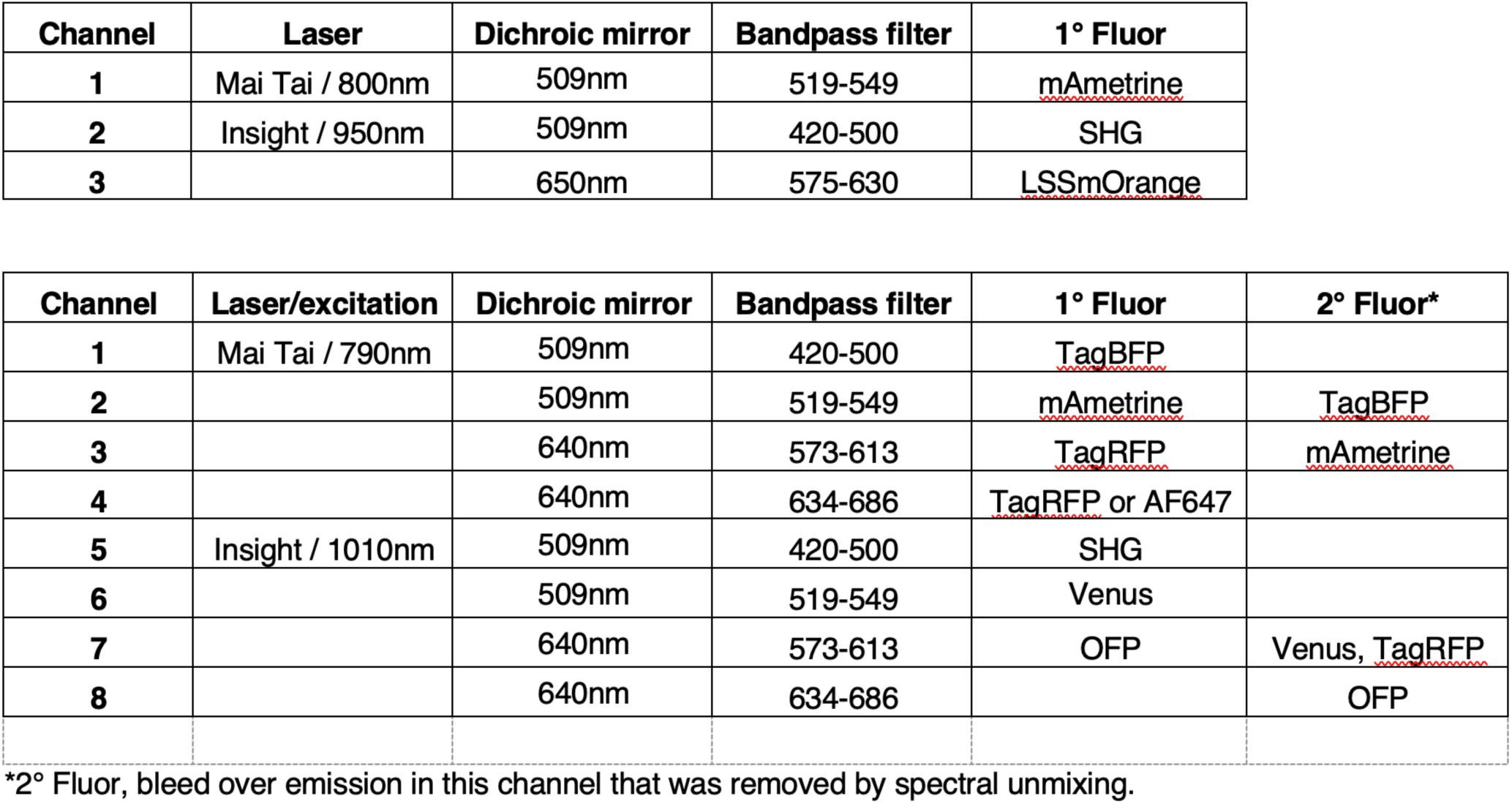
Two-photon microscope parameters.

**Supplemental Table S2.**
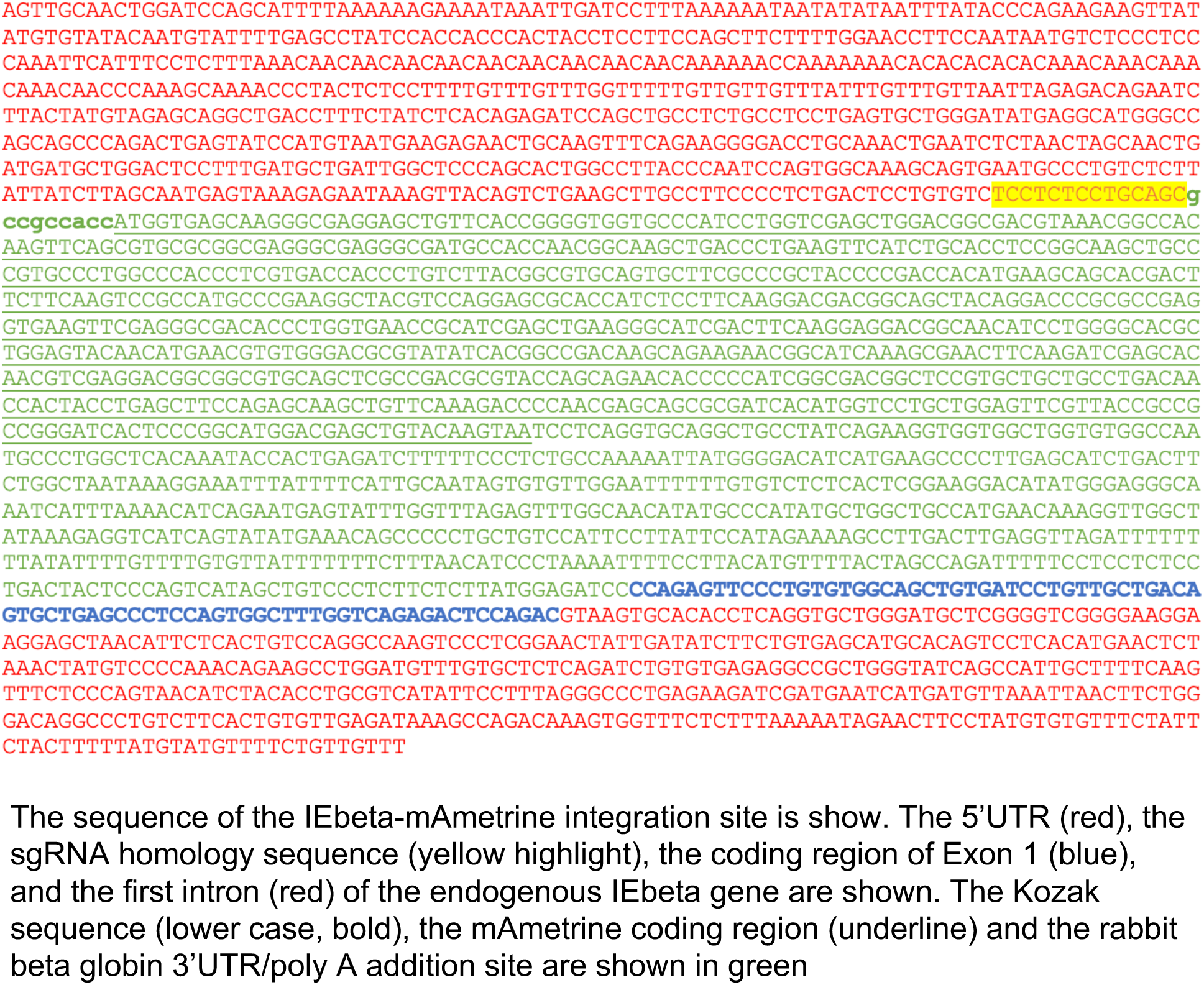
IEbeta-mAmetrine integration site.

**Supplemental Table 3.**
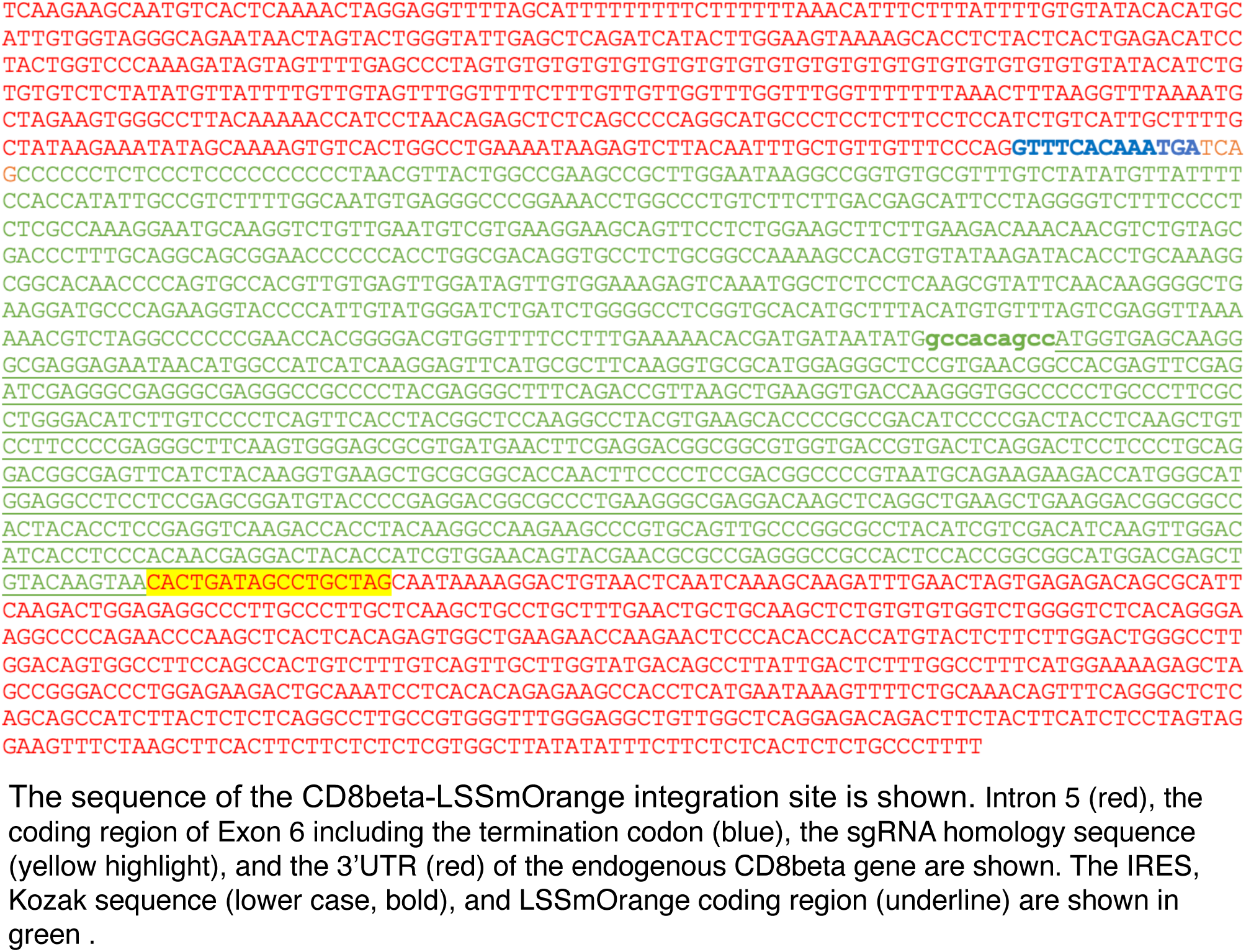
CD8-LSSmOrange integration site.

## References

Anderson KG, Mayer-Barber K, Sung H, Beura L, James BR, Taylor JJ, Qunaj L, Griffith TS, Vezys V, Barber DL, et al (2014) Intravascular staining for discrimination of vascular and tissue leukocytes. Nature Protocols 9: 209–222.

Bala N, McGurk AI, Zilch T, Rup AN, Carter EM, Leddon SA, Fowell DJ (2022) T cell activation niches-optimizing t cell effector function in inflamed and infected tissues. Immunol Rev 306: 164–180. doi:10.1111/imr.13047

Bannard O, McGowan SJ, Ersching J, Ishido S, Victora GD, Shin JS, Cyster JG (2016) Ubiquitin-mediated fluctuations in mhc class ii facilitate efficient germinal center b cell responses. J Exp Med 213: 993–1009. doi:10.1084/jem.20151682

Cheroutre H, Lambolez F (2008) Doubting the tcr coreceptor function of cd8alphaalpha. Immunity 28: 149–159. doi:10.1016/j.immuni.2008.01.005

Chieppa M, Rescigno M, Huang AY, Germain RN (2006) Dynamic imaging of dendritic cell extension into the small bowel lumen in response to epithelial cell tlr engagement. J Exp Med 203: 2841–2852. doi:10.1084/jem.20061884

Devine L, Kieffer LJ, Aitken V, Kavathas PB (2000) Human cd8b, but not mouse cd8b, can be expressed in the absence of cd8a as a bb homodimer. J Immunol 164: 833–838.

Di Pilato M, Kfuri-Rubens R, Pruessmann JN, Ozga AJ, Messemaker M, Cadilha BL, Sivakumar R, Cianciaruso C, Warner RD, Marangoni F, et al (2021) Cxcr6 positions cytotoxic t cells to receive critical survival signals in the tumor microenvironment. Cell 184: 4512–4530 e4522. doi:10.1016/j.cell.2021.07.015

Eidsmo L, Allan R, Caminschi I, van Rooijen N, Heath WR, Carbone FR (2009) Differential migration of epidermal and dermal dendritic cells during skin infection. J Immunol 182: 3165–3172. doi:10.4049/jimmunol.0802950

Ester M, Kriegel H-P, Sander J, Xu X (1996) A density-based algorithm for discovering clusters in large spatial databases with noise. Paper presented at Proceedings of the Second International Conference on Knowledge Discovery and Data Mining(KDD’96). AAAI Press;

Fung-Leung W-P, Kundig TM, Ngo K, Panakos J, De Sousa-Hitzler J, Wang E, Ohashi PS, Mak TW, Lau CY (1994) Reduced thymic maturation but normal effector function of cd8 + t cells in cd8beta gene-targeted mice. J Exp Med 180: 959–967.

Gangadharan D, Cheroutre H (2004) The cd8 isoform cd8alphaalpha is not a functional homologue of the tcr co-receptor cd8alphabeta. Curr Opin Immunol 16: 264–270. doi:10.1016/j.coi.2004.03.015

Gossa S, Nayak D, Zinselmeyer BH, McGavern DB (2014) Development of an immunologically tolerated combination of fluorescent proteins for in vivo two-photon imaging. Sci Rep 4: 6664. doi:10.1038/srep06664

Groom JR, Richmond J, Murooka TT, Sorensen EW, Sung JH, Bankert K, von Andrian UH, Moon JJ, Mempel TR, Luster AD (2012) Cxcr3 chemokine receptor-ligand interactions in the lymph node optimize cd4+ t helper 1 cell differentiation. Immunity 37: 1091–1103. doi:10.1016/j.immuni.2012.08.016

Hogquist KA, Jameson SC, Heath WR, Howard JL, Bevan MJ, Carbone FR (1994) T cell receptor antagonist peptides induce positive selection. Cell 76: 17–27.

Jansen CS, Prokhnevska N, Master VA, Sanda MG, Carlisle JW, Bilen MA, Cardenas M, Wilkinson S, Lake R, Sowalsky AG, et al (2019) An intra-tumoral niche maintains and differentiates stem-like cd8 t cells. Nature 576: 465–470. doi:10.1038/s41586-019-1836-5

Kissenpfennig A, Henri S, Dubois B, Laplace-Builhe C, Perrin P, Romani N, Tripp CH, Douillard P, Leserman L, Kaiserlian D, et al (2005) Dynamics and function of langerhans cells in vivo: Dermal dendritic cells colonize lymph node areas distinct from slower migrating langerhans cells. Immunity 22: 643–654. doi:10.1016/j.immuni.2005.04.004

Le Meur M, Gerlinger P, Benoist C, Mathis D (1985) Correcting an immune-response deficiency by creating ea gene transgenic mice. Nature 316: 38–42.

Leal JM, Huang JY, Kohli K, Stoltzfus C, Lyons-Cohen MR, Olin BE, Jr. MG, Gerner MY (2021) Innate cell microenvironments in lymph nodes shape the generation of t cell responses during type i inflammation. Science Immunology 6: eabb9435.

Li JL, Goh CC, Keeble JL, Qin JS, Roediger B, Jain R, Wang Y, Chew WK, Weninger W, Ng LG (2012) Intravital multiphoton imaging of immune responses in the mouse ear skin. Nat Protoc 7: 221–234. doi:10.1038/nprot.2011.438

Li JL, Goh CC, Ng LG (2018) Imaging of inflammatory responses in the mouse ear skin. Methods Mol Biol 1763: 87–107. doi:10.1007/978-1-4939-7762-8_9

Li S, Chen LX, Peng XH, Wang C, Qin BY, Tan D, Han CX, Yang H, Ren XN, Liu F, et al (2018) Overview of the reporter genes and reporter mouse models. Animal Model Exp Med 1: 29–35. doi:10.1002/ame2.12008

Lindquist RL, Shakhar G, Dudziak D, Wardemann H, Eisenreich T, Dustin ML, Nussenzweig MC (2004) Visualizing dendritic cell networks in vivo. Nat Immunol 5: 1243–1250. doi:10.1038/ni1139

Litingtung Y, Dahn RD, Li Y, Fallon JF, Chiang C (2002) Shh and gli3 are dispensable for limb skeleton formation but regulate digit number and identity. Nature 418: 979–983. doi:10.1038/nature01033

McRae TD, Oleksyn D, Miller J, Gao YR (2019) Robust blind spectral unmixing for fluorescence microscopy using unsupervised learning. PLoS One 14: e0225410. doi:10.1371/journal.pone.0225410

Mohan JF, Kohler RH, Hill JA, Weissleder R, Mathis D, Benoist C (2017) Imaging the emergence and natural progression of spontaneous autoimmune diabetes. Proc Natl Acad Sci U S A 114: E7776–E7785. doi:10.1073/pnas.1707381114

Natsuaki Y, Egawa G, Nakamizo S, Ono S, Hanakawa S, Okada T, Kusuba N, Otsuka A, Kitoh A, Honda T, et al (2014) Perivascular leukocyte clusters are essential for efficient activation of effector t cells in the skin. Nat Immunol 15: 1064–1069. doi:10.1038/ni.2992

Nesovic LD, Gonzalez Cruz PE, Rychener N, Wilks LR, Gill HS (2024) Standardizing the skin tape stripping method for sensitization and using it to create a mouse model of peanut allergy. Int J Pharm 662: 124479. doi:10.1016/j.ijpharm.2024.124479

Ng LG, Hsu A, Mandell MA, Roediger B, Hoeller C, Mrass P, Iparraguirre A, Cavanagh LL, Triccas JA, Beverley SM, et al (2008) Migratory dermal dendritic cells act as rapid sensors of protozoan parasites. PLoS Pathog 4: e1000222. doi:10.1371/journal.ppat.1000222

Pinkert CA, Wideral G, Cowing C, Heber-Katz E, Palmiter RD, Flavell RA, Brinster RL (1985) Tissue-specific, inducible and functional expression of the ead mhc classii gene intransgenicmice. The EMBO Journal 4: 2225–2230.

Prizant H, Patil N, Negatu S, Bala N, McGurk A, Leddon SA, Hughson A, McRae TD, Gao YR, Livingstone AM, et al (2021) Cxcl10(+) peripheral activation niches couple preferred sites of th1 entry with optimal apc encounter. Cell Rep 36: 109523. doi:10.1016/j.celrep.2021.109523

Reilly EC, Lambert Emo K, Buckley PM, Reilly NS, Smith I, Chaves FA, Yang H, Oakes PW, Topham DJ (2020) T(rm) integrins cd103 and cd49a differentially support adherence and motility after resolution of influenza virus infection. Proc Natl Acad Sci U S A 117: 12306–12314. doi:10.1073/pnas.1915681117

Romani N, Clausen BE, Stoitzner P (2010) Langerhans cells and more: Langerin-expressing dendritic cell subsets in the skin. Immunol Rev 234: 120–141. doi:10.1111/j.0105-2896.2009.00886.x

Sabino CP, Deana AM, Yoshimura TM, da Silva DF, Franca CM, Hamblin MR, Ribeiro MS (2016) The optical properties of mouse skin in the visible and near infrared spectral regions. J Photochem Photobiol B 160: 72–78. doi:10.1016/j.jphotobiol.2016.03.047

Srinivasan S, Zhu C, McShan AC (2024) Structure, function, and immunomodulation of the cd8 co-receptor. Front Immunol 15: 1412513. doi:10.3389/fimmu.2024.1412513

Tong PL, Roediger B, Kolesnikoff N, Biro M, Tay SS, Jain R, Shaw LE, Grimbaldeston MA, Weninger W (2015) The skin immune atlas: Three-dimensional analysis of cutaneous leukocyte subsets by multiphoton microscopy. J Invest Dermatol 135: 84–93. doi:10.1038/jid.2014.289

Ugur M, Mueller SN (2019) T cell and dendritic cell interactions in lymphoid organs: More than just being in the right place at the right time. Immunol Rev 289: 115–128. doi:10.1111/imr.12753

Victora GD, Nussenzweig MC (2022) Germinal centers. Annu Rev Immunol 40: 413–442. doi:10.1146/annurev-immunol-120419-022408

Yamamura K-i, Kikutani H, Folsom V, Clayton LK, Kimoto M, Akira S, Kashiwamura S-i, Tonegawa S, Kishimoto T (1985) Functional expression of a microinjected e! Gene in c57bl/6 transgenic mice. Nature 316: 67–69.

Yang SJ, Ahn S, Park CS, Holmes KL, Westrup J, Chang CH, Kim MG (2006) The quantitative assessment of mhc ii on thymic epithelium: Implications in cortical thymocyte development. Int Immunol 18: 729–739. doi:10.1093/intimm/dxl010

